# CRISPRi Gene Modulation and All-Optical Electrophysiology in Post-Differentiated Human iPSC-Cardiomyocytes

**DOI:** 10.1101/2023.05.07.539756

**Authors:** Julie L. Han, Yuli W. Heinson, Christianne J. Chua, Wei Liu, Emilia Entcheva

## Abstract

Uncovering gene-phenotype relationships can be enabled by precise gene modulation in human induced pluripotent stem-cell-derived cardiomyocytes (iPSC-CMs) and follow up phenotyping using scalable all-optical electrophysiology platforms. Such efforts towards human functional genomics can be aided by recent CRISPR-derived technologies for reversible gene inhibition or activation (CRISPRi/a). We set out to characterize the performance of CRISPRi in post-differentiated iPSC-CMs, targeting key cardiac ion channel genes, KCNH2, KCNJ2, and GJA1, and providing a multiparametric quantification of the effects on cardiac repolarization, stability of the resting membrane potential and conduction properties using all-optical tools. More potent CRISPRi effectors, e.g. Zim3, and optimized viral delivery led to improved performance on par with the use of CRISPRi iPSC lines. Confirmed mild yet specific phenotype changes when CRISPRi is deployed in non-dividing differentiated heart cells is an important step towards more holistic pre-clinical cardiotoxicity testing and for future therapeutic use in vivo.

## INTRODUCTION

The development of new integrative approaches to address human cardiac disease, which faithfully can predict *in vivo* phenotypes from gene expression profiling and from therapeutic gene modulation is highly desirable. CRISPR-derived tools for active gene inhibition or gene activation can empower such endeavors if successfully validated in human heart cells and tissues. CRISPR can target an exact genomic locus through designing specific short complementary nucleotide sequences. CRISPR/Cas9 gene editing relies on the precise nuclease action to generate double-strand breaks (DSB) and the activation of DNA repair pathways. The efficiency of the mechanisms to repair the damaged DNA can be variable dependent on cell type^1^. Homology-directed repair (HDR) is an error free repair mechanism, but is restricted to S and G2 phases of the cell cycle, and therefore, this pathway is not suitable to repair the genome in non-dividing cells, such as adult cardiomyocytes^1, 2^. DSB-involving CRISPR gene editing is particularly challenging in induced pluripotent stem cells, iPSC, and derivatives as these cells show high cytotoxicity in response to the DSB cuts^3–5^.

Over the last decade, multiple efforts have led to the development of newer CRISPR-derived tools to avoid DSBs for safer and more efficient gene control^6^. Some of these methods, e.g. prime editing^7^ and base editing^8, 9^, that use mutant or inactive version of the Cas9 enzyme and avoid the need for DSB and HDR, have accelerated so quickly as they are now entering clinical trials. Interference and activation CRISPR (CRISPRi/a) belong to this class of approaches. They use a deactivated Cas9 (dCas9) fused to an effector (a repressor^10, 11^ or activator^12^) to achieve specific and reversible gene modulation. The most common version of CRISPRi uses the Krüppel-associated box (KRAB) repression domain (dCas9-KRAB) for superior transcriptional knockdown without cytotoxic effects when compared to using active Cas9. Mandegar et al.^13^ reported an inducible CRISPRi platform in human iPSCs coupled with RNAseq that was not only reversible but also outperformed CRISPR with an active Cas9. They showed utility in studying genes involved in cardiac cell differentiation as well as cardiac repolarization (KCNH2), documenting phenotypic responses, i.e. prolongation of the action potential in iPSC-CMs. Limpitikul et al.^14^ applied CRISPRi to target calmodulin mutations associated with long QT syndrome and corrected the action potential duration in patient-derived iPSC-CMs. In the few cardiac studies that have deployed CRISPRi/a, the efforts have been focused on creating stable dCas9-expressing iPSC lines^13, 15, 16^. While dCas9 did not prevent follow up differentiation of the iPSCs into cardiac cells of different lineage, no comprehensive comparison was done to reveal potential effects on the iPSC-CM phenotype. There is a motivation to be able to deploy the CRISPRi/a gene modulation approaches in any patient-derived iPSC-CMs for large-scale screens of loss-of-function and gain-of function perturbations using pooled or arrayed libraries of guide RNAs, gRNAs^3, 17^, without the need to create stable dCas9 iPSC lines, including the use of standard commercial iPSC-CMs to avoid variability associated with in house differentiation^18, 19^. Better understanding of the direct use of CRISPRi in non-dividing human cardiomyocytes can also be informative for future in vivo deployment in the post-natal human heart.

In this study, we combined CRISPRi gene modulation and scalable all-optical electrophysiology to link gene perturbation to complex phenotypes in commercially available post-differentiated iPSC-CMs. We targeted three major ion channels implicated in cardiac electromechanical function: *KCNH2*, *KCNJ2*, and *GJA1* and quantitatively linked multiparametric functional responses to protein levels and mRNA levels using an experimental pipeline (**Figure 1**). A thorough characterization of the use of CRISPRi in human iPSC-CMs is presented, including assessment of potential side effects and variability of functional responses in different iPSC-CM lines even for the same gene knockdown efficiency. We also tested for the first time a newer CRISPRi effector – Zim3^20^ – to control gene transcription in human iPSC-CMs. Optimization of these tools for effective gene inhibition/activation, can speed up *in vivo* therapeutic use. Overall, the reported approaches and results can inform future gene-function investigations to better understand the molecular underpinnings of cardiac electromechanics in health and disease, and in a patient-specific manner, to aid the development of safer (non-cardiotoxic) new therapeutics.

**Figure 1.**
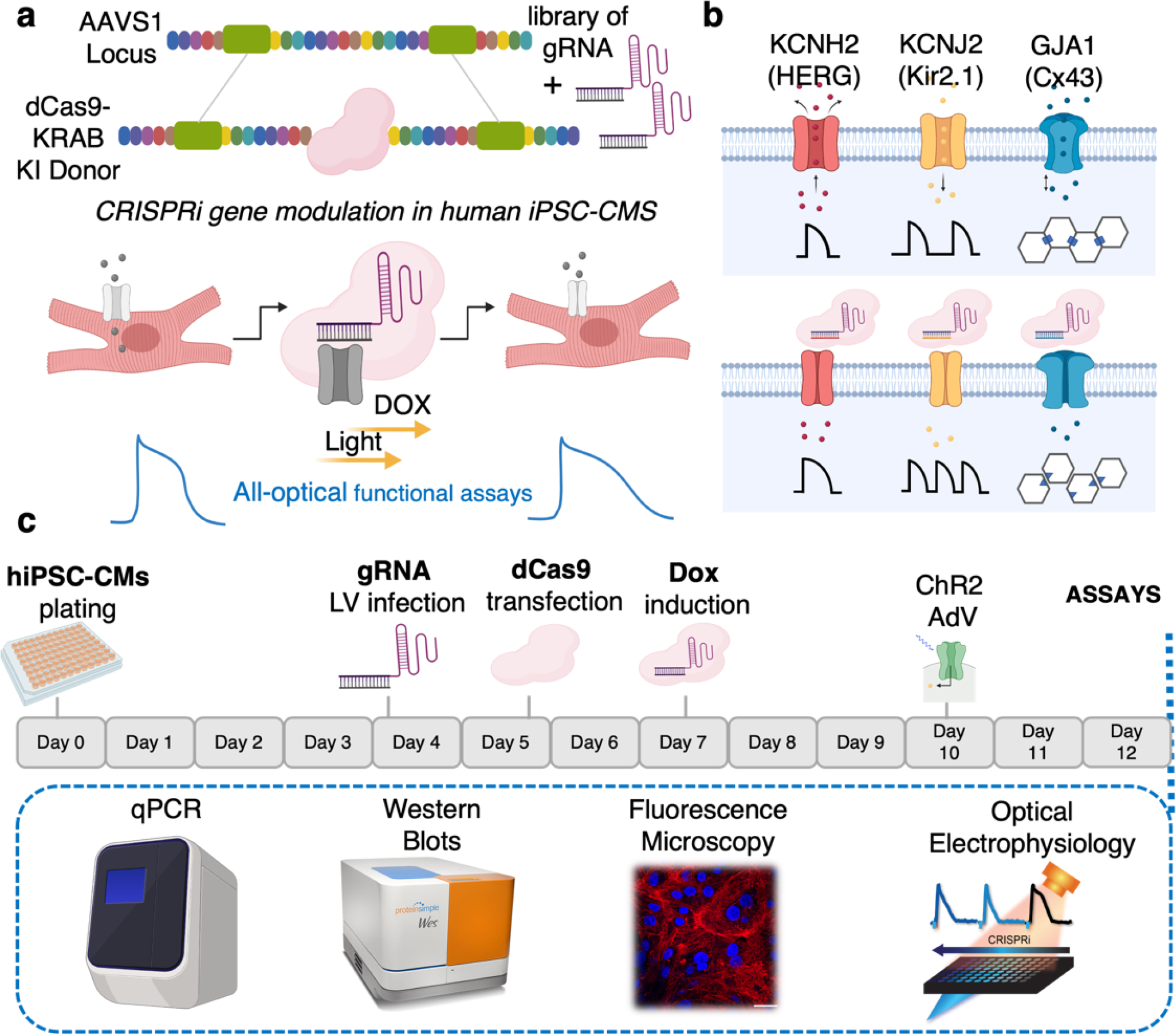
Scalable platform for CRISPRi gene modulation in post-differentiated human iPSC-CMs followed by functional assessment using optical electrophysiology and other assays – schematic and timeline. This platform utilizing 96-well format has the potential to be scaled to test libraries of gRNA perturbing genes specific to studying cardiac electrophysiology, disease modelling, and toward drug discovery. a, dCas9-KRAB is inserted into the *AAVS1* locus site in post-differentiated human iPSC-CMs and, with the addition of a library of gRNAs, we can conduct full all-optical functional assays in tandem with CRISPR Cas9 mediated gene modulation. b, As a proof of principle, we focus on characterizing the functional outputs of *KCNH2*, *KCNJ2*, and *GJA1* knockdown. c, Diagram showing the experimental pipeline (see Online Methods). Post-differentiated hiPSC-CM are transduced with gRNA-expressing lentivirus, transfected with a Dox-inducible dCas9-KRAB, and treated with Dox (2 μM) for 5-days prior to functional, mRNA, and protein analyses. In all microscopic studies, cells were also transduced with ChR2, an optogenetic actuator.

## RESULTS

A feasibility CRISPRi study was performed in post-differentiated iPSC-CMs targeting key genes important in cardiac electrophysiology. Comprehensive analysis using all-optical electrophysiology and a pipeline enabling correlative analysis of functional and molecular data in the same samples helped quantify the CRISPRi gene modulation in this *in vitro* model.

### Time-Dependent Control of dCas9-KRAB and its Tracing by Fluorescence in Live Cells

Optimal control of gene expression features high activity in the presence of an effector, but low background in its absence. Inducible systems, such as Doxycycline-induced (Dox), allow temporal control of expression. In this study, the inducible (Tet-on) dCas9-KRAB-mCherry was inserted into the AAVS1 locus in hiPSC-CMs (**Figure 1**) as described in the Methods. The dCas9-KRAB expression post-Dox treatment was quantified both using terminal Western blot (WB) assays and by mCherry live tracking. After Dox treatment, mCherry expression increased with some delay, reaching about 50% of its maximum 24h after initial introduction (**Figure 2a**). WB quantification demonstrated that the CRISPRi system begins to express dCas9 protein earlier than reported by mCherry and it reached peak expression at about 24h post initial Dox treatment (**Figure 2b**). Relatively sustained expression was seen both at the fluorescent level and protein level for 48h during Dox treatment (**Figure 2a, b**). The data shows that dCas9-KRAB protein and its fluorescent tag can be controlled with the Tet-on promotor, supporting studies that rely on precise time-dependent gene inhibition.

**Figure 2.**
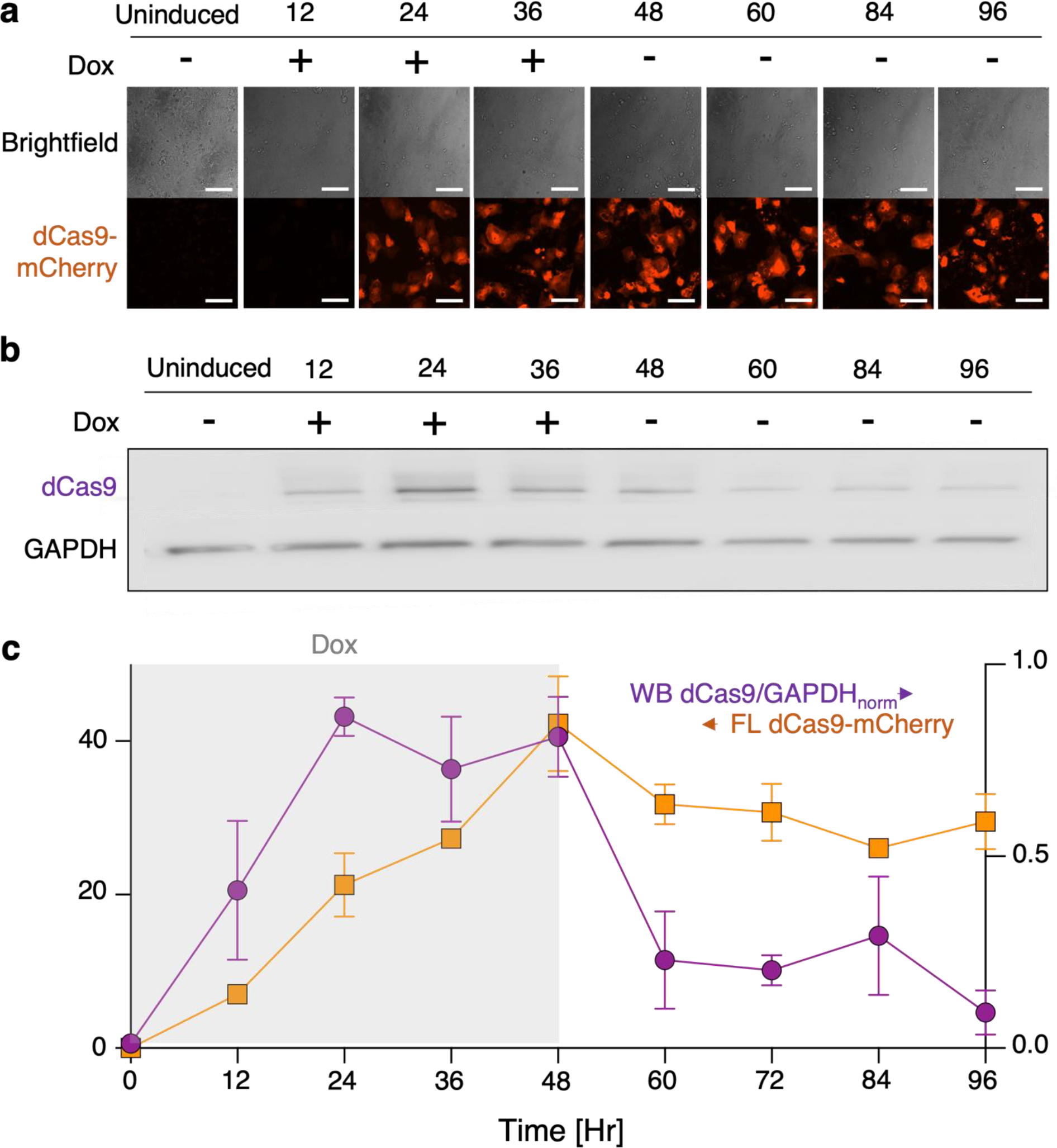
Temporal control of dCas9-KRAB expression by Dox in the AAVS1 locus in iPSC-CM. a, Tracing dCas9-mCherry expression in live hiPSC-CMs by fluorescence (FL); Dox (2 μM) was introduced at Hour 0 and removed at Hour 48; n≥3 biologically independent samples per timepoint. **b,** Quantification of dCas9-KRAB protein expression at respective time points by Western Blot (WB); n≥2 biologically independent samples per timepoint. **c,** Comparison of the kinetics of FL- and WB-determined dCas9- KRAB expression; Dox (2 μM) application is shown in gray shading. Data are presented as mean ± SEM. Scale bar 100 *μ*m.

The dCas9 expression, as reported by WB and by live mCherry fluorescence, differed both in the on and especially in the off kinetics as Dox was applied or removed. Upon Dox removal, over 80% of dCas9 degradation took place in the first 12h after removal with complete recovery within 48h, based on WB results (**Figure 2b, c**). In contrast, sustained mCherry fluorescence was seen 48h after removal of Dox (**Figure 2a, c**). The discrepancy in kinetics between the two quantification methods alerts us about some limitations of live fluorescence reporting. Contributing to this is likely mCherry’s resistance to proteolysis and accumulation in the lysosomes^21^, as well as slower maturation compared to other fluorescent reporters^21, 22^. Overall, our results corroborate the importance of developing inducible Cas9 systems with higher temporal resolution to optimize gene modulation despite limitations of the lengthy time scales of transcription and translation^13, 23^.

### Potential Side Effects of the CRISPRi Components on Cardiac Electrophysiology

In a complex deployment of a multiparameter system for gene control, it is important to quantify potential side effects of various treatments and components on the output variables of interest. Dox has been reported to have minimal eukaryotic off-target effects as its main use is bacteria-specific. With Dox treatment of post-differentiated hiPSC-CMs, we sought to characterize any molecular or functional electrophysiological effects beyond overt cytotoxicity. Dox application alone had no statistically significant effects on key electrophysiological parameters used in this study – action potential duration (APD) and calcium transient duration (CTD) under spontaneous or paced conditions; it also did not alter spontaneous frequency of activity after 5 days of treatment (**Suppl. Figure 1a;** left**, 1a;** bottom right). Furthermore, we sought out to determine if potential Dox-induced off-target effects can alter the transcription of our genes of interest. No significant difference was found at the mRNA level for KCHN2, KCNJ2, and GJA1 upon Dox treatment alone (**Suppl. Figure 1b**). There were also no significant changes at the protein level – for Kir2.1 and Cx43 as quantified by standard WB and WES (Protein Simple) (**Suppl. Figure 1c**). The lack of side effects from Dox application led to our decision to perform most comparisons of CRISPRi gene modulation in this study between +/-Dox treatment.

Separately, we sought to determine if insertion of dCas9-KRAB in the AAVS1 locus has no off-target effects on our ion channels of interest. Indeed, no significant difference was seen for Kir2.1 and Cx43 protein levels in dCas9-KRAB expressing cells when compared to eGFP and mCherry control groups as quantified by both standard WB and WES (ProteinSimple)(**Suppl. Figure 1d**).

Finally, we took a closer look at the gRNA lentiviral delivery. Polybrene is a cationic polymer that is commonly used to increase the transduction efficiencies of retroviruses in cell culture, including for lentiviral vectors. It is a relatively non-toxic polymer that acts by neutralizing the charge repulsion between virions and sialic acid on the cell surface and has been reported to have minimal effects on cells even during prolonged use in culture^24^. However, in our experience, when polybrene was used to introduce the gRNA packaged in lentivirus, we observed about a 20-40% prolongation in spontaneously recorded APD80 (**Suppl. Figure 2**). Separately, paradoxically, polybrene appeared to decrease the transfection efficiency of dCas9-KRAB in cells after their transduction with gRNA (**Suppl. Figure 2a**) despite previous studies showing that polybrene increased gene transfer efficiency by lipofection^25^. Due to the off-target effects we observed in functionality and with no advantageous contributions of using polybrene in our system, we excluded it as a reagent for the delivery of the gRNAs and settled on the proposed methodology (**Figure 1**).

### CRISPRi of KCNH2 in hiPSC-CMs and its Functional Consequences on Repolarization

The KCNH2 gene encodes hERG, the voltage sensitive potassium (K^+^) channel protein, which mediates the rapid delayed rectifier K^+^ current (IKr)^26^. IKr is a major contributor to the repolarization phase of the cardiac action potentials, effectively influencing the action potential duration, APD, and the QT interval observed in electrocardiograms. Loss-of-function mutations or drug-induced reduction in hERG lead to APD prolongation experimentally and QT interval prolongation clinically, therefore increasing the risk for fatal arrhythmias^27^. Preclinical testing of all new drugs brought to market assess for hERG block because of key role for hERG in drug-induced arrhythmia^28^.

To deploy CRISPRi for hERG manipulation, we tested multiple gRNA’s for KCNH2, designed around the transcription start site (TSS) where it is known to be most effective; ultimately we found that a previously published gRNA (175) used with human iPSC-CMs^13^ exhibited the most efficient knockdown and therefore used it in subsequent functional experiments, **Figure 3a, Suppl. Figure 3**. In our expression system (**Figure 1c**) transforming post-differentiated human iPSC-cardiomyocytes, we saw 40% knockdown of KCNH2 at the mRNA level, **Figure 3c, e.** These levels are slightly reduced compared to differentiated cardiomyocytes using dCas9-KRAB iPS cell line. Mandegar et al. observed > 95% suppression in the stable dCas9-KRAB iPS lines but only 60% suppression the in differentiated iPSC-CMs ^13^.

**Figure 3.**
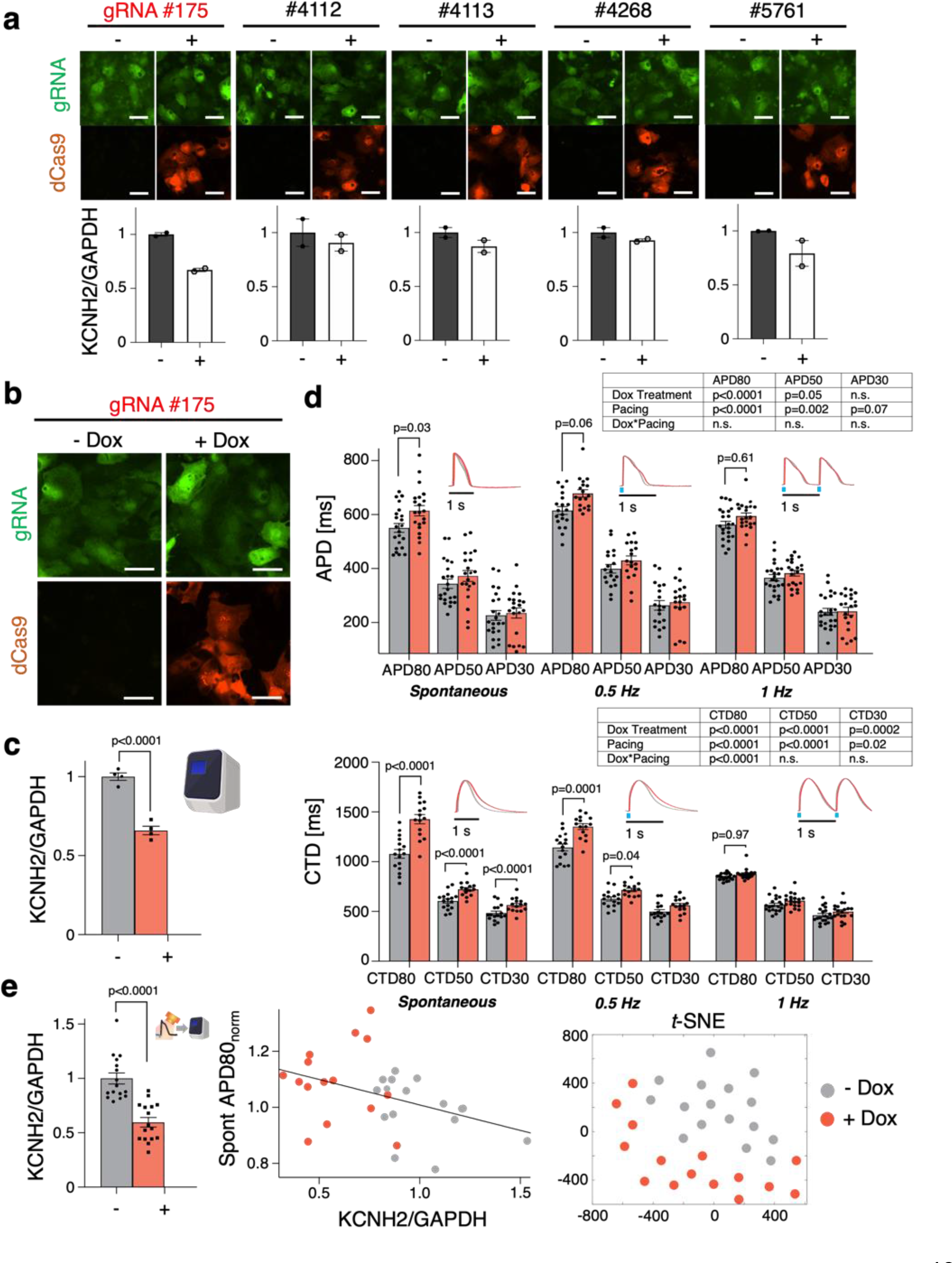
**Knockdown of *KCNH2* by CRISPRi results in functional repolarization changes**. **a,** Validation in identifying most efficient gRNA for *KCNH2* knockdown (n=2 biologically independent samples, n=3 technical replicates per gRNA tested). **b,** Fluorescence expression of dCas9-mCherry and gRNA 175 tagged with eGFP upon 5-days of Dox (2 μM) treatment. **c,** qPCR of CRISPRi knockdown of KCNH2 with gRNA 175 in hiPSC-CMs (n=4 biologically independent samples, n=3 technical replicates; unpaired t-test). **d,** Functional changes in APD and CTD in Dox (2 μM) treated samples expressing Dox- inducible dCas9-KRAB and gRNA targeting *KCNH2* (n≥18 biologically independent samples and n≥15 biologically independent samples, respectively; two-way ANOVA). **e,** qPCR analysis of *KCNH2* knockdown post-optical electrophysiology (n=16 biologically independent samples, n=3 technical replicates; unpaired t-test). Plot correlating the effects of relative *KCNH2* mRNA to spontaneous APD80 with a linear best of fit line shown (n=15 biologically independent samples). *t*-NSE dimension reduction plot generated encompassing full set of functional and genetic data (19 measured parameters) collected from KCNH2 CRISPRi knockdown samples (n=15 biologically independent samples). Data are presented as mean ± SEM. Scale bar 50 *μ*m.

Upon CRISPRi knockdown, we saw about 12% APD80 prolongation and about 40% CTD80 prolongation during spontaneous oscillations (**Figure 3d**, top and bottom, respectively). Slower pacing rates uncovered larger APD prolongation compared to faster pacing. The mild effects of CRISPRi on APD are consistent with results obtained using dCas9-KRAB iPS lines and their differentiation into cardiomyocytes^13^. After functional assessment with all-optical electrophysiology, we were able to apply qPCR analysis in the same samples to quantify the mRNA levels of KCNH2 knockdown and observed that 40% knockdown was conserved upon running the qPCR post-labeling with optical sensors for voltage and calcium and using an optogenetic actuator (**Figure 3e**, left). Having molecular and functional data from the same samples enabled correlative analysis. We visualized the data for the two experimental groups: -Dox and +Dox. (**Figure 3e**, middle). As our all-optical electrophysiology assay and pipeline processing outputs comprehensive multiparameter data, we used a dimension-reduction approach and visualized the two groups in a *t*-SNE plot. The 19 different parameters: relative mRNA quantity, spontaneous frequency, various APD and CTD features under spontaneous and paced (0.5 Hz and 1 Hz) conditions – were projected onto a latent space via *t*-SNE (**Figure 3e**, right). Despite the mild effects of CRISPRi KCNH2 knockdown on APD prolongation, group separation was seen in this projection that integrates multiparameter information.

### Electrophysiological Phenotype of CRISPRi Knockdown of KCNJ2 in hiPSC-CMs and Effects of Cell Density

KCNJ2 gene encodes for the potassium inward rectifier channel Kir2.1 protein (I_K1_ current) - a critical contributor to the stability of the resting membrane potential and to cardiac excitability^29–31^. Gain-of-function or loss-of-function mutations in KCNJ2 cause sudden cardiac death syndromes, and reduction in I_K1_, is a contributing factor to arrhythmogenesis in failing human hearts^29, 30^. The levels of expression of KCNJ2 and the functionality of I_K1_ have been recognized as a key metric for electrical maturity of hiPSC-CMs and constitute an important consideration in drug safety testing^31–34^. One of the manifestations of deficient I_K1_ current is an increase in spontaneous oscillations due to the more depolarized resting membrane potential. In this study we targeted KCNJ2 with CRISPRi, testing a panel of 5 gRNAs designed around the TSS start site to isolate the best candidate to conduct functional studies. Of the tested gRNAs, KCNJ2 ERA2 was deemed the most effective, with up to 70% knockdown, **Figure 4a**.

**Figure 4.**
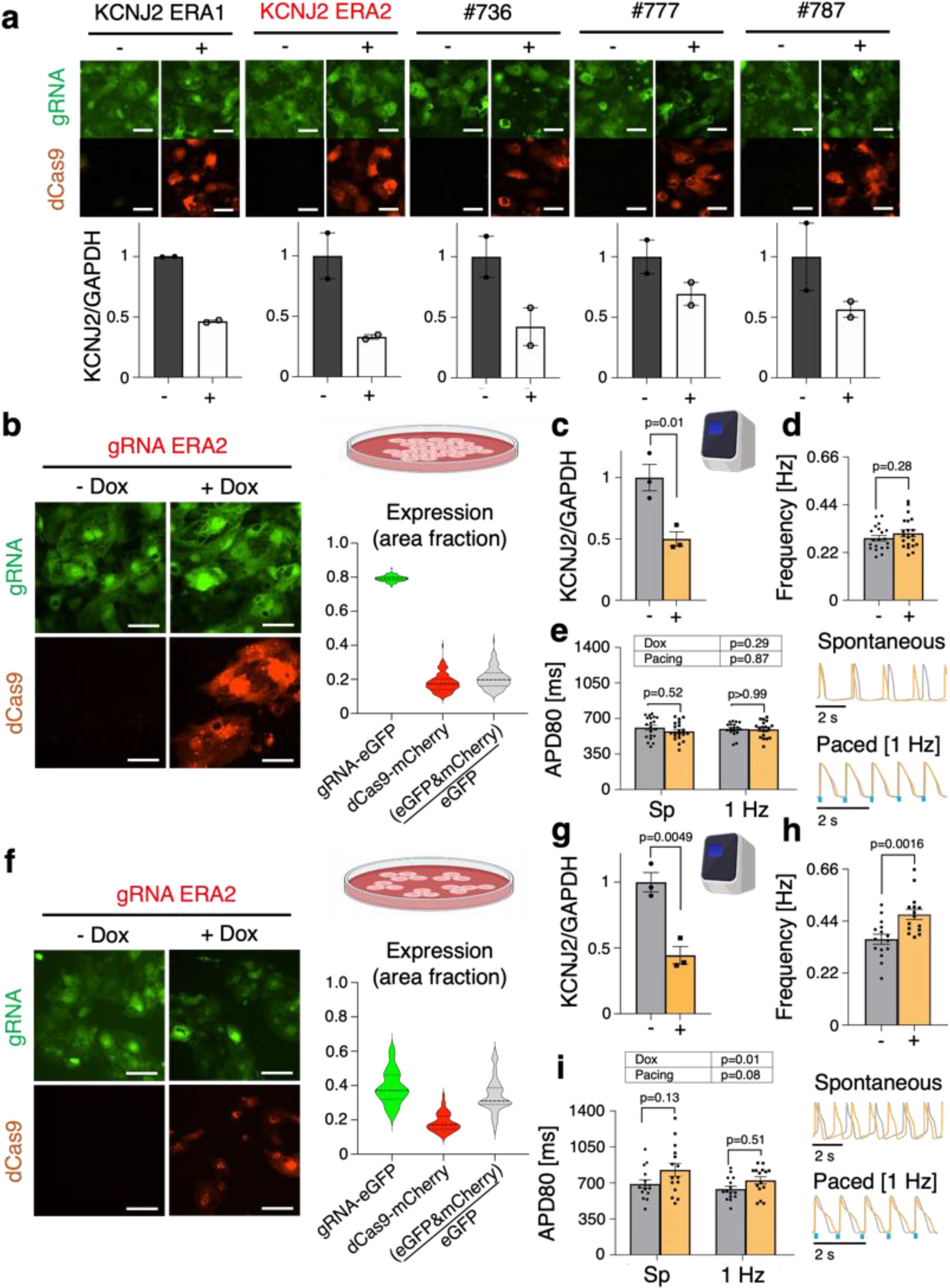
The electrophysiological effects of CRISPRi knockdown of *KCNJ2* on spontaneous rates are enhanced in reduced cell densities. a, Validation in identifying most efficient gRNA for *KCNJ2* knockdown (n=2 biologically independent samples, n=3 technical replicates per gRNA tested). (**b – e**), Data for high cell density plating groups. **b,** Fluorescence expression of dCas9-mCherry and gRNA ERA2 tagged with eGFP upon 5-days of Dox (2 μM) treatment in high cell density plated groups. Expression ratios were calculated based on pixel area of mCherry and eGFP (see Online Methods); (n=126). **c,** qPCR of CRISPRi knockdown of *KCNJ2* with gRNA ERA2 in hiPSC-CMs (n=3 biologically independent samples, n=3 technical replicates; unpaired t-test). **d,** Frequency changes in Dox (2 μM) treated cardiomyocytes expressing Dox-inducible dCas9-KRAB and gRNA targeting *KCNJ2* (n=20 biologically independent samples; unpaired t-test). **e,** APD80 changes in Dox (2 μM) treated groups from both spontaneous (Sp) and paced data sets (n>18 biologically independent samples; two-way ANOVA). (**f – i**), Data for reduced/medium cell density plating groups. **f,** Fluorescence expression of dCas9-mCherry and gRNA-eGFP upon 5-days of Dox (2 μM) treatment in medium cell density plated groups. Expression ratios were calculated based on pixel area of mCherry and eGFP (see Online Methods); (n=48). **g,** qPCR of CRISPRi knockdown of KCNJ2 in medium density plated hiPSC-CMs (n=3 biologically independent samples, n=3 technical replicates; unpaired t-test). **h,** Frequency changes in Dox-inducible dCas9-KRAB and gRNA targeting *KCNJ2* in medium density plated hiPSC-CMs (n=15 biologically independent samples; unpaired t-test). **i,** APD80 changes in Dox (2 μM) treated groups from both Sp and paced data sets (n=15 biologically independent samples; two-way ANOVA). Data are presented as mean ± SEM. Scale bar 50 *μ*m.

In high density plated hiPSC-CMs (our typical 50,000 cells per 96-well), despite about 50% knockdown of KCNJ2 (**Figure 4c**), there was only a slight increase in the spontaneous beat frequency (**Figure 4d**) and no change in APD80 for both spontaneous and paced data sets (**Figure 4e**). In highly-coupled syncytium and incomplete spatially “spotty” CRISPRi (see **Figure 4b** and **Suppl. Figure 4**), stronger KCNJ2 from the unaffected cells can smooth out the localized knockdown, thereby reducing the effectiveness of the perturbation. We speculated that the effects of the KCNJ2 knockdown would be enhanced if the coupling is slightly reduced. Cells were plated at three-fold lower density and the experiments were repeated. At this seeding density, gRNA ERA2 exhibited comparable knockdown efficiency at about 55% knockdown (**Figure 4g**). As expected, more pronounced functional differences were revealed, with about 30% increase in spontaneous frequency in the CRISPRi samples compared to control (**Figure 4h**). Additionally, cells showed APD80 prolongation in spontaneous and paced conditions (**Figure 4i**, bottom left). The relatively mild effects in high density groups and more pronounced effects in lower density groups, reinforces our and others’ previous findings of the importance of syncytial conditions when studying the electrophysiology of hiPSC-CMs^35–37^.

### CRISPRi of GJA1 and Effects on Conduction in hiPSC-CMs

Gap junctional communication is critical in the heart for maintaining coordinated excitation and coordinated contraction^38, 39^. Cell-cell coupling in the heart is mediated by cardiac gap junction channels, from the connexins (Cx) family, of which Cx43 is the predominant ventricular isoform. Altered expression and distribution of connexins is a hallmark in the pro-arrhythmic remodeling associated with structural heart disease^40^. Here we sought to characterize CRISPRi knockdown of GJA1 gene which encodes for Cx43. Five sgRNAs were tested, of which GJA1 1218 exhibited the highest suppression at about 50% knockdown at the mRNA level and was used for further functional experiments, **Figure 5a, c**. At the protein level, gRNA 1218 exhibited about 25% knockdown as quantified by WES (ProteinSimple), **Figure 5d**. Functionally, there was overall slowing of conduction velocity at 0.5 Hz, 0.75 Hz, and 1 Hz pacing frequencies with most pronounced effects at the higher pacing frequencies, as the system is stressed, **Figure 5e**. Our pipeline for optical electrophysiology experiments followed by mRNA or protein analysis in the same samples allowed the generation of correlation plots with CV at 1Hz and the respective normalized mRNA levels of GJA1/GAPDH and Cx43/GAPDH protein levels, **Figure 5f**.

**Figure 5.**
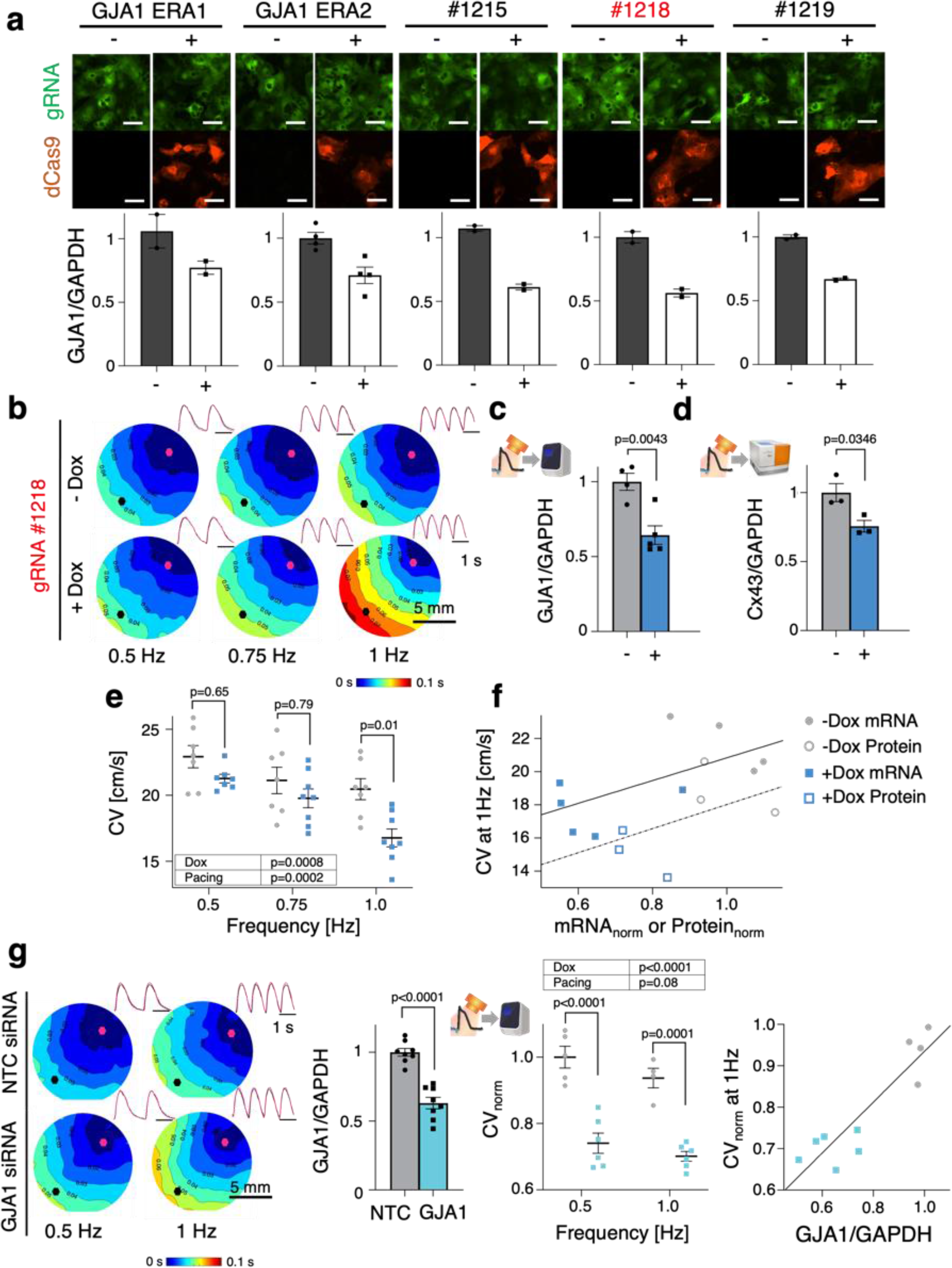
CRISPRi of *GJA1* reduces conduction velocity, as shown by optical mapping**. a,** Validation in identifying most efficient gRNA for *GJA1* knockdown (n>2 biologically independent samples, n=3 technical replicates per gRNA tested). **b,** Activation maps of hiPSC-CMs knocked down by CRISPRi and gRNA 1218 at 0.5Hz, 0.75Hz and 1 Hz pacing frequencies; representative images (n>7 biologically independent samples) **c,** qPCR of CRISPRi knockdown of *GJA1* with gRNA 1218 post-optical electrophysiology (n>4 biologically independent samples, n=3 technical replicates; unpaired t-test). **d,** WES (ProteinSimple) protein quantification of *GJA1* knockdown by CRISPRi in hiPSC-CMs post-optical electrophysiology (n=3 biologically independent samples, unpaired t-test). **e,** Conduction velocity changes in Dox (2 μM) treated samples at 0.5 Hz, 0.75 Hz, and 1 Hz pacing (n>7 biologically independent samples, two-way ANOVA). **f,** Plot correlating relative *GJA1* mRNA levels and relative Cx43 protein levels with CV at 1 Hz pacing (n>3 biologically independent samples). **g,** Activation maps of *GJA1* knockdown of hiPSC-CMs with siRNA at 0.5 Hz and 1 Hz pacing frequencies; representative images (n>4 biologically independent samples). qPCR of *GJA1* knockdown by siRNA post-optical electrophysiology (n=8 biologically independent samples, n=3 technical replicates; unpaired t-test). Conductional velocity changes in siRNA knockdown samples at 0.5 Hz and 1 Hz pacing (n>4 biologically independent samples; two-way ANOVA). Plot correlating relative *GJA1* mRNA levels with CV at 1 second (n>4 biologically independent samples). Data are presented as mean ± S.E.M. Scale bar 50 *μ*m.

To determine how the more established approach of siRNA knockdown compares to CRISPRi, we conducted additional experiments using siRNA targeting GJA1 (**Figure 5g**). Similarly, to the CRISPRi knockdowns, we saw about 40% knockdown of GJA1 at the mRNA level (**Figure 5g**). However, the CV decrease at both at 0.5 Hz and 1 Hz pacing was more pronounced with siRNA. Correlation plots also could be generated with CV at 1Hz related to the respective mRNA levels of GJA1/GAPDH to better visualize the two experimental groups (**Figure 5g**). Despite CRISPRi and siRNA having similar knockdown efficiencies, conduction velocity data resulted in more pronounced effects using siRNA targeting GJA1. This may be explained by a more uniform global effect of siRNA compared to the spatially “spotty” CRISPRi knockdown, as in the case of KCNJ2. RNAi, which utilizes endogenous protein machinery already present in mammalian cells^41^, may exhibit some advantages over dCas9 if the delivery of Cas9 constructs is sub-optimal.

### Application of CRISPRi Across Different hiPSC-CM Cell Lines

To test if the findings from this study, obtained in the commercially-available and optimized iCell^2^ ventricular cell line (from an 18yr. old female) are generalizable, we explored using CRISPRi in additional hiPSC-CM cell lines for KCHN2 and GJA1 knockdown. The gRNA #175 exhibited comparable knockdown efficiencies to iCell^2^ in a cell line from a male aged 50-59yr at about 40%, **Figure 6b**. Interestingly, despite similar knockdown efficiencies, the male hiPSC-CMs exhibited more pronounced APD responses with about 30% APD80 prolongation in spontaneous recordings, **Figure 6a, right**. After all-optical electrophysiology, the 40% KCNH2 knockdown efficiency was generally preserved and we were able to construct correlation plots between mRNA and functional metrics for the -Dox and +Dox group, **Figure 6c**. Additionally, we used CRISPRi to knockdown GJA1 in the same cell line and observed slowing of conduction velocity at 0.5 Hz pacing in the Dox-treated group, **Figure 6d**. From these samples, we conducted qPCR to quantify GJA1 mRNA and observed about 45% knockdown, similar to the iCell^2^ line. Correlation plots visualized the links between CV at 0.5 Hz pacing and the respective GJA1/GAPDH mRNA levels, **Figure 6e**. Additional CRISPRi tests for the same two genes were performed in a third cell line from a female aged 50-59yr. The functional results were milder for this cell line, which overall had significantly slower conduction velocity and shorter action potentials at baseline compared to the other two cell lines, **Suppl. Figure 5**. Particularly for KCNH2 knockdown, the functional APD prolongation was not present, **Suppl. Figure 5a, right**. These data emphasize the need for considering population responses and personalized testing.

**Figure 6.**
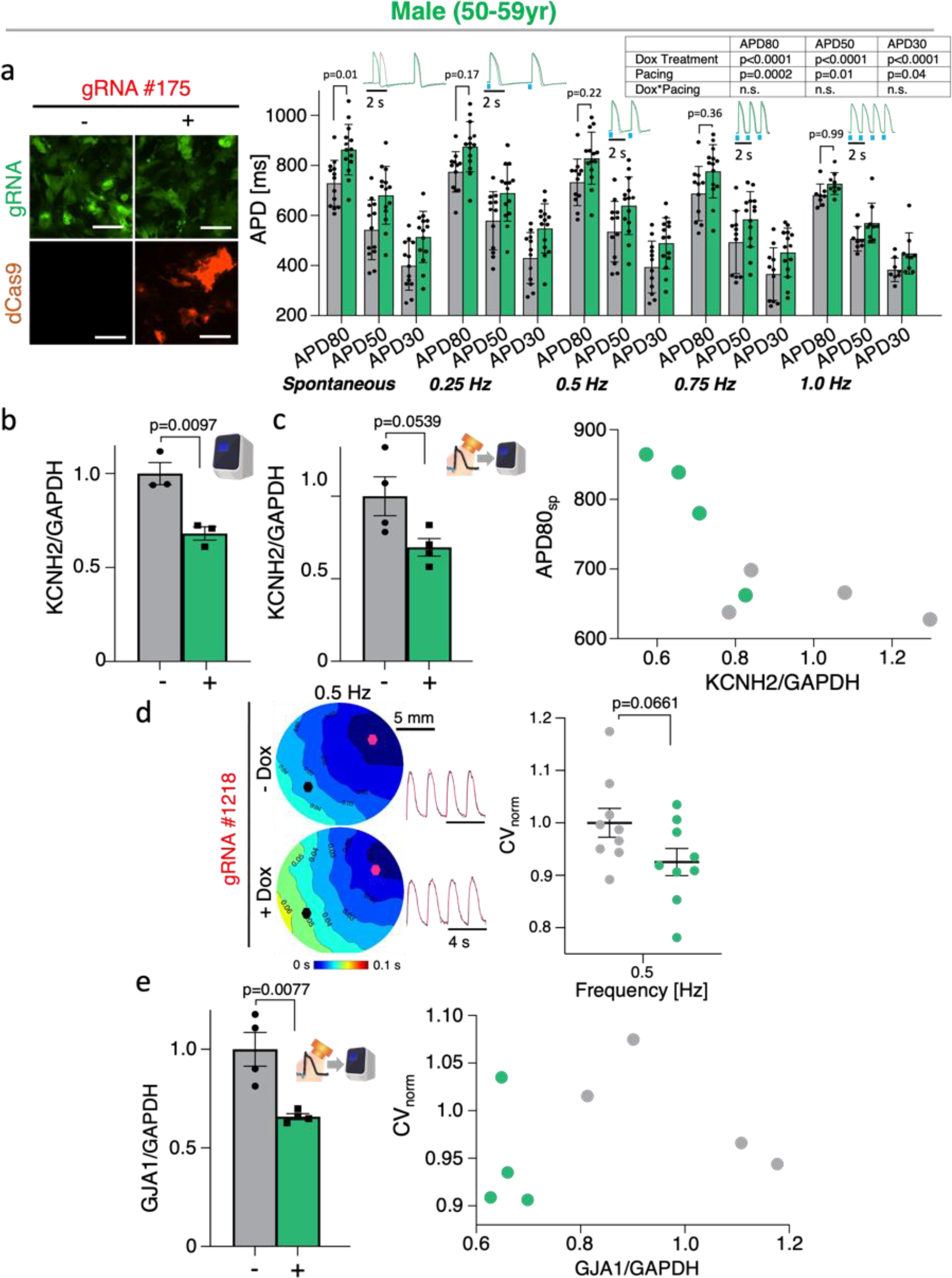
**Translation of CRISPRi tools to alternative hiPSC-CM cell lines**. **a**, Fluorescence expression of dCas9-mCherry and gRNA 175 tagged with eGFP upon 5-days of Dox treatment in male (50-59yr) hiPSC-CMs; representative images. Functional changes in APD in Dox (2 μM) treated samples expressing Dox-inducible dCas9-KRAB and gRNA targeting *KCNH2* (n>8 biologically independent samples; two-way ANOVA). **b**, qPCR of CRISPRi knockdown of *KCNH2* with gRNA 175 in male hiPSC- CMs (n=3 biologically independent samples, n=3 technical replicates; unpaired t-test). **c,** qPCR analysis of *KCNH2* knockdown post-optical electrophysiology (n=4 biologically independent samples, n=3 technical replicates; unpaired t-test). Plot correlating the effects of relative *KCNH2* mRNA to spontaneous APD80 (n=4 biologically independent samples). **d,** Activation maps of male hiPSC-CMs knocked down by CRISPRi and gRNA 1218 at 0.5Hz pacing; representative images (n=4 biologically independent samples; unpaired t-test). Conduction velocity changes in Dox (2 μM) treated samples at 0.5 Hz pacing (n=9 biologically independent samples; unpaired t-test). **e,** qPCR of CRISPRi knockdown of *GJA1* with gRNA 1218 post-optical electrophysiology (n=4 biologically independent samples, n=3 technical replicates; unpaired t-test). Plot correlating relative *GJA1* mRNA levels with CV at 0.5 Hz pacing (n=4 biologically independent samples). Data are presented as mean ± SEM. Scale bar 100 *μ*m.

### HT Studies for Human Cardiac Functional Genomics using Improved CRISPRi Effector Zim3

When combined with CRISPR gene modulation, high-throughput platforms for all-optical electrophysiology^42–46^, enabled by optogenetics^47^, can open the door to human functional genomics studies in the cardiac field. Using 96-well plate-reader all-optical platform, developed in our lab as derivative of recent work^48, 49^, **Figure 7a**, we obtained synchronous, multi-well measurements of KCNH2 knockdown allowing us to compare the Dox-inducible dCas9-KRAB, siRNA, and a recent more efficient CRISPRi system, dCas9-KRAB-Zim3^20^, using the Zinc Finger Imprinter 3 gene as part of the effector complex. We developed an adenoviral vector to more uniformly deliver the dCas9-KRAB-Zim3 construct. Successful expression of the dCas9-KRAB-Zim3 was confirmed in the hiPSC-CMs, **Figure 7b.** Of the three methods of KCNH2 knockdown, we observed that the recently developed dCas9-KRAB-Zim3 (at MOI 1000) induced the most significant APD prolongation in the hiPSC-CMs upon KCNH2 knockdown. Separately, in these paced datasets, dCas9-KRAB-Zim3 had no effect on CTD, **Figure 7d**. Using our pipeline, we combined the relative KCNH2 mRNA data obtained post-optical electrophysiology (**Figure 7e**) and the APD80 at 0.5 Hz pacing data from the same samples to conduct correlative analysis, **Figure 7f**. In additional experiments, even a two-fold lower MOI (500) of dCas9-KRAB-Zim3, induced a significant knockdown of KCNH2, **Suppl. Figure 6b**, and concomitant but milder prolongation of APD80 at 0.5 Hz pacing, **Suppl. Figure 6c**. The effect on APD was significant across pacing frequencies, **Suppl. Figure 6d, left**, without a significant change of CTD80 at the respective pacing frequencies, **Suppl. Figure 6d, right**. The relative KCNH2 mRNA data and the APD80 at 0.5 Hz pacing were used to conduct correlative analysis, **Suppl. Figure 6e**. Interestingly, we observed an increase in relative KCNH2/GAPDH levels in dCas9-KRAB-Zim3 infected hiPSC-CMs compared to non-transformed control post-labelling with optical sensors for voltage and calcium, using ChR2 for optogenetic actuation, **Figure 7e**. For this reason, all comparisons were between scramble gRNA and gRNA against a target of interest.

**Figure 7.**
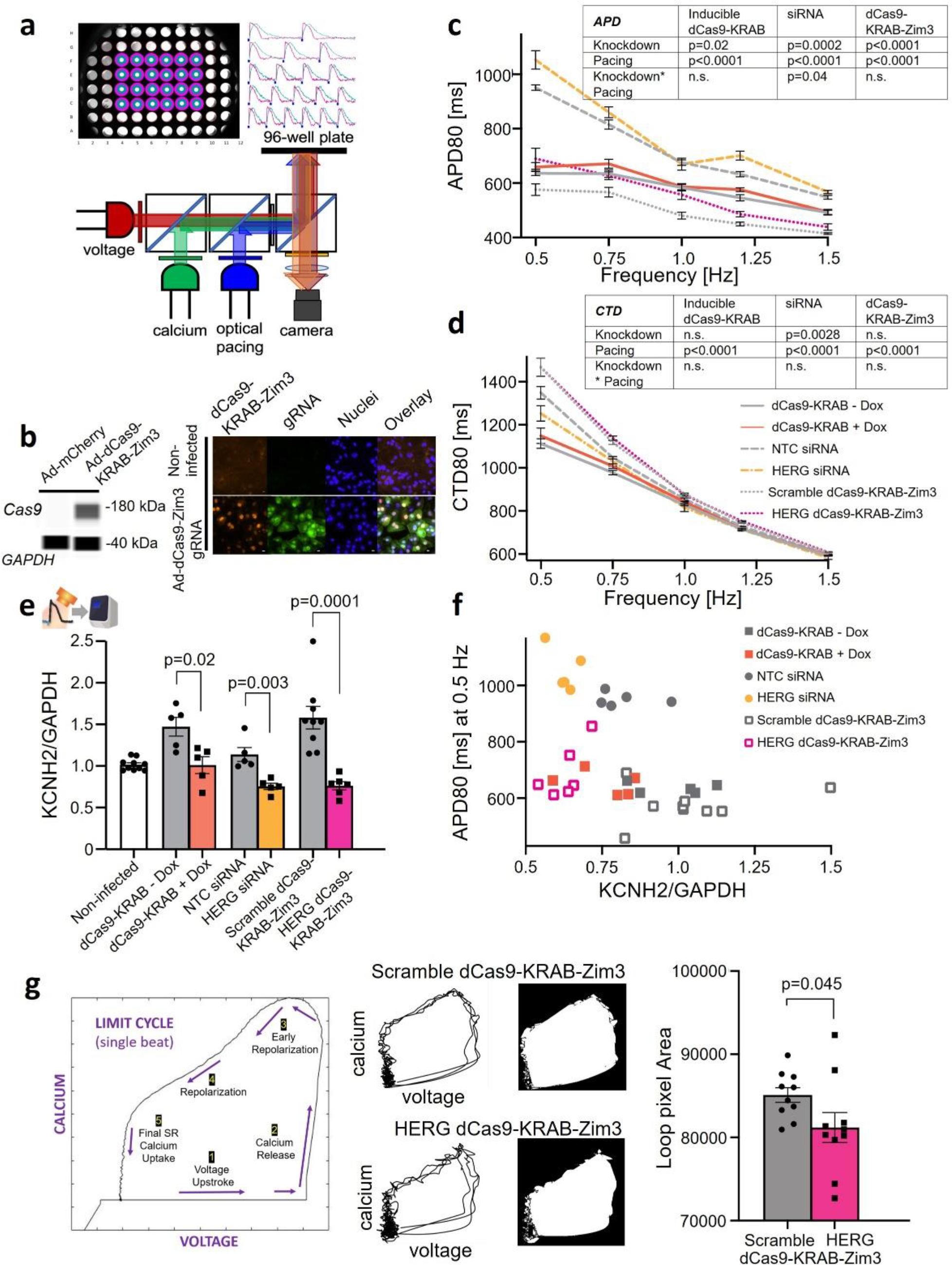
**High-throughput (HT) studies for synchronous, multi-well measurements with improved Zim3 CRISPRi**. **a**, Schematic illustrating optical system capable of simultaneous optical pacing and voltage and calcium imaging of an entire 96-well plate. HT, all-optical electrophysiology studies were conducted at 35°C. **b**, WES (ProteinSimple) of hiPSC-CMs expressing dCas9-Zim3; representative images (WES, n=4 biologically independent samples). **c,** Restitution plot of APD80 upon KCNH2 knockdown using inducible dCas9-KRAB (Gen1) (n=5 biologically independent samples; two-way ANOVA), siRNA (n=4-5 biologically independent samples; two-way ANOVA), and dCas9-KRAB-Zim3 (n=6-12 biologically independent samples; two-way ANOVA). **d,** Restitution plot of CTD80 upon KCNH2 knockdown using inducible dCas9-KRAB (Gen1) (n=5 biologically independent samples; two-way ANOVA), siRNA (n=5 biologically independent samples; two-way ANOVA), and dCas9-KRAB-Zim3 (n=6- 11 biologically independent samples; two-way ANOVA). **e,** qPCR of KCNH2 knockdown using inducible dCas9-KRAB (Gen1) (n=5 biologically independent samples, n=3 technical replicates; unpaired t-test), siRNA (n=5 biologically independent samples, n=3 technical replicates; unpaired t-test), and dCas9- KRAB-Zim3 (n=6-9 biologically independent samples, n=4 technical replicates; unpaired t-test). **f,** Plot correlating relative *KCNH2* mRNA levels with APD80 at 0.5 Hz pacing. **g,** Example time-embedded voltage-calcium loop with indicated phases. Upon knockdown of KCNH2 by dCas9-KRAB-Zim3, there are changes in both the shape and limit cycle pixel area (n=10 biologically independent samples per group; unpaired t-test). Data are presented as mean ± SEM. Scale bar 10 *μ*m.

Embedded-time limit cycles can be used to capture the relational dynamics of variables in a system, **Figure 7g, left**. Such loops were generated using the voltage and calcium traces of KCNH2 knockdown by dCas9-KRAB-Zim3, **Figure 7g, middle**. Upon reduction of I_Kr_, APD is prolonged and it extends to a later phase of the calcium transient, so that the last vertical segment of final calcium uptake by the SR becomes shorter and the loop slightly collapses. Such decrease in the loop pixel area is documented here for hiPSC-CMs with KCNH2 knockdown compared to control, **Figure 7g, right**, indicative of a slower repolarization as expected. Similar to the latent space projection (t-SNE), this approach integrates multiparameter information about a phenotype induced by a specific gene modulation.

## DISCUSSION

In this study, we developed a system for CRISPRi gene modulation in commercially available post-differentiated cardiomyocytes. We demonstrated its capacity for transcriptional control in such differentiated cells, targeting genes of key importance for cardiac electrophysiology. Furthermore, we implemented a pipeline for sequential functional characterization using all-optical electrophysiology and follow-up quantification of gene expression in the same samples. This approach allows for unique correlative and potentially causative explorations of gene-phenotype relationships.

For iPS and iPSC-CMs, CRISPRi can be viewed as a superior option for gene inhibition (knockdown) because of lower cytotoxicity, potentially higher efficiency, and a possibility for time-resolved, reversible action^6^, compared to standard CRISPR knockout. Cytotoxicity of standard CRISPR knockout comes from error-prone DNA repair of the double-strand breaks^4, 5^. Dose and temporal control of gene modulation are desirable features to avoid off-target effects, chromosomal translocations, and genotoxicity^23^. Unlike CRISPR knockout, time-resolved CRISPRi allows the study of essential genes and their role in cellular functions. In our Dox-inducible CRISPRi system, we observed minimal side effects (**Suppl. Figure 1**). The choice of post-differentiated cardiomyocytes in this study was with the intent to avoid potential interferences with the differentiation process, to avoid potential TetO promoter silencing (Dox induction)^13, 50^ during the Wnt-signaling modulation required for cardiac differentiation, and to provide testing in conditions that are a step closer to *in vivo* deployment for gene modulation.

Overall, we observed specific, but mild effects from CRISPRi knockdown of the three key cardiac ion channels studied: HERG, Kir2.1, and Cx43. There were challenges in finding a highly efficient gRNA targeting the KCNH2 gene in post-differentiated cardiomyocytes, despite testing multiple gRNAs selected close to the TTS (**Figure 3a, Suppl. Figure 3**). We were able to achieve only about 40% knockdown efficiency (**Figure 3c, e**). Mandegar et al. report they observed a higher knockdown efficiency using the same gRNA in iPSCs compared to iPSC-CMs differentiated from the stable dCas9 iPSC lines^13^. The sub- optimal efficiency of the dCas9-KRAB in our study may be due in part to using transfection in most experiments (**Suppl. Figure 4a**). In contrast, lentiviral delivery of the gRNAs yielded over 90% expression (**Suppl. Figure 4a**). Previous studies using CRISPR/Cas9 *in vivo* have shown genetic mosaicism in both edited and non-edited cells^51^. Yet, despite observations of inconsistent expression, phenotypic changes were reported^51, 52^. In our case, the functional effects (prolongation of APD) were frequency-dependent - differences were more pronounced for slower pacing rates and were abolished at faster rates – an effect known as “reverse rate dependence”, which most hERG-blocking drugs exhibit^53^. These findings suggest the importance of optimizing each respective model system and the need for better delivery systems both *in vivo* and *in vitro*.

KCNJ2 is a critical gene in stabilizing the resting membrane potential and preventing spontaneous oscillations in ventricular cells, thus considered a determinant of iPSC-CM maturity. To uncover the expected functional effects of CRISPRi on KCNJ2, we had to use hiPSC-CMs grown at reduced cell density (**Figure 4f, Suppl. Figure 4b**). The importance of seeding density and syncytium formation on I_K1_ in hiPSC-CM has been discussed before^36, 54^. In both the high density and medium density groups, we achieved similar knockdown efficiency (**Figure 4c, g**). In conditions of non-uniform, “spotty” suppression of KCNJ2, the electrotonic coupling in dense cultures can smooth out local effects on the membrane potential. In reduced cell density conditions, the CRISPRi modulation of KCNJ2 yielded expected increase in spontaneous oscillations compared to control.

Similar to the syncytium-dependent effects of suppressing KCNJ2, we also explored the effects of CRISPRi on gap junctional genes, e.g. GJA1, and its role in conduction properties of multicellular preparations. Frequency-dependent responses revealed that GJA1 suppression is more pronounced at higher pacing rates, upon critical slowing of conduction velocity, **Figure 5**. Very few studies have directly linked mRNA and protein levels of gap junctional proteins to conduction velocity changes^48^. Here the combination of CRISPRi, all-optical electrophysiology and our pipeline for processing the samples permitted such correlative analysis, **Figure 5f-g**. Using standard siRNA resulted in more pronounced functional effects of GJA1 knockdown (**Figure 5g**) as well as KCNH2 knockdown (**Figure 7c, d**). A likely explanation is the better targeting - although transfection was used for both, the single-component small siRNAs likely affected more cells compared to the co-expression of both dCas9 and the gRNA delivered separately in our case.

For a subset of experiments, we prepared an adenoviral vector using a recently developed improved effector for CRISPRi - dCas9-KRAB-Zim3^20^. Combined with our high-throughput all-optical electrophysiology platform – a full 96-well imager, derived from recently built systems^48, 49^, we were able to compare the performance of siRNAs, dCas9-KRAB and dCas9-KRAB-Zim3. As expected, the knockdown efficiency was improved, likely due to the viral delivery, **Figure 7e**. The Zim3 effector also showed the highest specificity and efficiency in suppressing KCNH2 (resulting in frequency dependent APD prolongation), **Figure 7**. Additionally, the Zim3 construct is smaller and therefore packaging for in vivo delivery may be easier. The highly parallel all-optical platforms, as the one deployed here, can facilitate the testing and optimization of new effectors for CRISPRi/a or other gene modulation tools, by providing a comprehensive functional readout. For visualization and classification in phenotyping, dimension-reduction approaches, e.g. t-SNE and voltage-calcium time-embedded loops, can be useful, as illustrated in **Figures 3e** and **7g**.

For translatability of gene modulation tools to personalized medicine applications, it is important to consider inter-subject differences. Comprehensive studies of over 700 hiPSC lines from over 300 donors have observed variable molecular signatures – both donor-specific and of unknown sources^18, 19^. Varied responses to pharmacological perturbations were found in studies as part of the CiPA initiative using donor-specific cell lines^55, 56^. There have been reports for the differential efficiency of CRISPRi/a in hiPSC-derived neurons from different donors^57^. Therefore, we tested the inducible CRISPRi tools in two additional commercial post-differentiated human iPSC-CM lines - a male (**Figure 6**) and an additional female hiPSC-CM line, both 50-59yr old (**Suppl. Figure 5**). Despite similar knockdown efficiency of the KCNH2 gene in all three cell lines (**Figure 3c, Figure 6b, Suppl. Figure 5b**), varied functional responses were observed. The male line showed the most significant differences in APD prolongation upon KCNH2 knockdown (**Figure 6a**), when compared to both female lines (**Figure 3d, Suppl. Figure 5a**). The additional female line (50-59yr) showed no statistically significant difference in APD prolongation (**Suppl. Figure 5a**). Overall, the functional responses of this diversity line were inferior at baseline – slower conduction velocity and triangular action potentials. Due to variability of different lines, as well as potential immaturity of some lines compared to others, parallelization and scalable testing is desirable to better capture population responses to identical gene modulation.

Human iPSC-CMs are an invaluable tool to study cardiac development and cardiac diseases, modelling monogenic and complex genetic diseases and interacting factors. And with the continuous developments of newer CRISPR-derived tools for gene modulation^20, 58–60^, with better tunability, more robust activity, and easier packaging, this experimental model can empower human functional genomics. The impact for cardiac research can come from the combination of three scalable technologies: human iPSCs, all-optical electrophysiology and CRISPRi/a screens, particularly arrayed screens, where the functional impact of each gene perturbation is studied^6^. Such comprehensive analysis can yield valuable information for building gene regulatory networks, correlative and causal models of cardiac gene regulation, and help drug development, cardiotoxicity testing and cell optimization for regenerative medicine applications.

## METHODS

### Culture and Maintenance of hiPSC-CMs

iCell Cardiomyocytes^2^ (Cat. C1016, Donor 01434, Fujifilm/Cellular Dynamics), diversity panel female iCell Cardiomyocytes (Cat. IPSC-CM-1X-01064, Fujifilm/Cellular Dynamics) and male iCell Cardiomyocytes (Cat. IPSC-CM-1X-01063, Fujifilm/Cellular Dynamics) were used. **Suppl. Table 1** lists details about the cell lines. Cardiomyocytes were cultured on fibronectin (Cat. 356009, Corning) coated (50μg/mL) glass-bottom 96-well plates (Cat. P96-1-N, Cellvis) at 5.0 x 10^4^ hiPSC-CMs cells per well and maintained according to manufacturer’s protocol. In some experiments 2.7 x 10^5^ hiPSC-CMs were plated on fibronectin-coated 35mm dish with a 14mm glass-bottom well for macroscopic optical mapping for conduction assessment. All cell types were mycoplasma tested and cultured at 37°C and 5% CO_2_.

### Sequence and Time Course of Experimentation

**Figure 1c** illustrates the sequence of perturbations and the assays performed. To enable correlative analysis, in most experiments, samples were processed through a pipeline: first functionally characterized using all-optical electrophysiology and then lysed to perform qPCR or Western Blot on the same samples and relate function to gene and protein expression.

### CRISPRi Gene Modulation in hiPSC-CMs

#### gRNA Design and Lentivirus Production

Guide RNAs (gRNA) were designed to target around the transcription start site (TSS) using CRISPR-ERA (Stanford) or VectorBuilder’s automated sgRNA design software. **Suppl. Table 2** lists all gRNA oligo sequences used in this study. All sgRNA’s were cloned into a lentiviral vector expressing an eGFP fluorescent reporter and commercially packaged into lentiviruses by VectorBuilder.

#### CRISPRi with dCas9-KRAB

For the main experiments, doxycycline (Dox)-inducible CRISPRi system (with KRAB effector domain) was used, where dCas9-KRAB-mCherry was inserted in the AAVS1 locus using Talens. Specifically, five days after plating, hiPSC-CMs were transfected with 0.0625μg AAVS1-Talen-L, a gift from Danwei Huangfu (Cat. 59025, Addgene), 0.0625μg AAVS1-Talen-R, a gift from Danwei Huangfu (Cat. 59026, Addgene), and 0.125μg pAAVS1-Ndi-CRISPRi (dCas9-KRAB, Gen1 – inducible system), a gift from Bruce Conklin^13^ (Cat. 73497, Addgene) per 5 x 10^4^ hiPSC-CMs using Lipofectamine 3000 transfection agent (Cat. L3000001, ThermoFisher) following manufacturer’s instructions. 2 μM Dox (Cat. D9891, Sigma-Aldrich) was introduced 2 days post-transfection and maintained for the totality of the culture.

For experiments characterizing Dox effects on hiPSC-CMs (**Suppl. Figure 1a, b**), cells were treated with 2 μM Dox initially at day 7 post-plating and Dox was supplemented for 5 days before lysing the cells for mRNA and protein quantification, following the timeline of Dox treatment during typical CRISPRi knockdown studies. For experiments characterizing the effects of dCas9-KRAB on ion channels (**Suppl. Figure 1d**), hiPSC-CMs were transfected with 0.0625ug AAVS1-Talen-L, 0.0625μg AAVS1-Talen-R, and 0.125μg of either: AAVS1-Pur-CAG-mCherry, a gift from Su-Chun Zhang (Cat. 80946, Addgene), AAV-CAGGS-eGFP, a gift from Rudolf Jaenisch (Cat. 22212, Addgene), or pAAVS1-NC-CRISPRi (dCas9- KRAB, Gen3 – constitutive expression), a gift from Bruce Conklin^13^ (Cat. 73499, Addgene) per 5 x 10^4^ hiPSC-CMs 5 days post-plating with Lipofectamine 3000. Cells were collected for protein analysis 2 days post-transfection.

#### CRISPRi with dCas9-KRAB-Zim3

For dCas9-KRAB-Zim3 studies, we used an adenovirus Ad-dCas9-mCherry-Zim3 (VectorBuilder), based on plasmid pHR-UCOE-SFFV-dCas9-mCherry-ZIM3-KRAB, a gift from Mikko Taipale^20^ (Cat. 154473, Addgene). To remain consistent with the Dox-inducible dCas9-KRAB methodology, (**Figure 1c**), hiPSC-CMs were transduced with gRNA lentivirus on Day 4 post-plating and Ad-dCas9-mCherry-Zim3 5 days post-plating. hiPSC-CMs were transduced with Ad-Zim3 at MOI 1000 in (**Figure 7**) and MOI 500 (**Suppl. Figure 6**). Cells were labeled for functional measurements and collected for mRNA analysis on Day 10 to allow for 5 days of sustained dCas9-Zim3 expression.

### siRNA Gene Suppression

In a subset of macroscopic optical mapping experiments, we used siRNA for gene suppression of GJA1. Cells were transfected with 270ng of esiRNA Human GJA1 (Cat. EHU105621, Millipore Sigma) or siRNA Universal Negative Control (Cat. SIC001, Millipore Sigma) per 2.7 x 10^5^ hiPSC-CMs 5 days post plating using Mission® siRNA Transfection Reagent (Cat. S1452, Millipore Sigma) following manufacturer’s instructions. Cells were assayed and collected for qPCR 4 days post-transfection. For HT studies, cells were transfected with 50ng of esiRNA Human KCNH2 (Cat. EHU080951, Milipore Sigma) or siRNA Universal Negative Control per 5.0 x 10^4^ hiPSC-CMs using Mission® siRNA Transfection Reagent, following manufacturer’s instructions. Cells were assayed and collected for qPCR 4 days post-transfection.

### All-Optical Functional Measurements

For CRISPRi knockdown studies, dCas9-KRAB transfection was done on day five post-plating, and Dox was introduced 2 days later. Then Channelrhodopsin-2 (ChR2) was expressed using adenoviral transduction with Ad-CMV-hChR2(H134R)-eYFP (Vector Biolabs) to cells to undergo all-optical functional assays on Day 12, (**Figure 1c**).

All microscopic functional assays were conducted using OptoDyCE^44, 45^ on hiPSC-CM monolayers plated in fibronectin-coated 96-well format in Tyrode’s Solution (in mM): NaCl, 135; MgCl_2_, 1; KCl, 5.4; CaCl_2_, 1.8; NaH_2_PO_4_, 0.33; glucose, 5.1; and HEPES, 5; adjusted to pH 7.4 with NaOH, at room temperature. The OptoDyCE system uses a single camera (iXon Ultra 897 EMCCD; Andor Technology Ltd., Belfast, UK) and temporal multiplexing to record optically both voltage and calcium transients in response to optogenetic stimulation. It is built around an inverted microscope (Nikon Ti2 with programmable stage) and uses NI Elements for the recordings and control. TTL-programmable LEDs allow optical actuation (5ms pulses) with a 470nm, 650 mW LED (Thorlabs; Newton, NJ), controlled with an LED driver (Thorlabs). Temporal multiplexing of green (530nm) and red LED (660nm) (Thorlabs) allows voltage and calcium recordings. Calcium was recorded using Rhod-4AM at 10uM (AAT Bioquest, Sunnyvale, CA) with fluorescent excitation/emission peaks at 530 nm and 605 nm respectively, and voltage was tracked using a near-infrared voltage dye BeRST1 at 1uM (from Evan W. Miller, University of California, Berkeley) with fluorescent excitation/emission at 660 nm and 700nm long pass, respectively^45^. Signals were filtered and analyzed using an automated custom software to quantify relevant parameters^44, 45^. Typical records included trains of spontaneous activity as well as optically paced activity at several frequencies, most often at 0.5Hz and 1Hz.

For imaging macroscopic waves over a field of view of about 1.2x1.5cm, hiPSC-CMs were plated onto 35mm dishes with 14mm glass-bottom and genetic modifications were scaled by cell number. The widefield optical wave mapping was conducted with custom built systems, as described^48, 49^. High-speed machine vision CMOS cameras, (Basler, Ahrensburg Germany) were used in conjunction with oblique LED trans-illumination (green for calcium and red – for voltage, as described above). Either electrical (5ms pulses via bipolar platinum electrodes) or optical initiation (10ms pulses using 470nm at 0.28 mW/mm^2^). Signals were filtered and analyzed with an automated custom software to construct activation maps and compute conduction velocity^48^. Such macroscopic wave imaging was used in **Figures 5, 6** and **Suppl. Figure 5**. In recent developments, the stand-alone macroscopic system^49^ has been modified to image a whole 96-well plate in epi-illumination mode, using the same camera. In some cases, time-embedded voltage-calcium loops were constructed from the dual measurements under optical pacing. The area of such loops was calculated after the 2D plots were converted to images, images were binarized, loop area was filled in and pixels within the area counted. High-throughput (HT) data are shown in **Figure 7** and **Suppl. Figure 6**.

### qRT-PCR Analysis

Cells were harvested for total RNA and quantified using the POWER SYBR^TM^ Cells-To-C_T_ Kit (Cat. 4402955, ThermoFisher Scientific) following the manufacturer’s protocol. qPCR analysis was performed on the QuantStudio 3 Real-Time PCR System (ThermoFisher Scientific) to detect each target gene (Table S2) and analyzed using QuantStudio^TM^ Design & Analysis Software (ThermoFisher). Linearity of the response and primer efficiency was confirmed using serial dilution of cDNA. All samples were normalized to housekeeping gene GAPDH and results were expressed as the relative mRNA normalized to control using the standard ΔΔCt method^61^.

### Protein Quantification

#### Standard Western Blot

Cells were harvested for total protein using a Qproteome Mammalian Protein Prep Kit (Qiagen). Total protein was quantified using a Pierce^TM^ BCA Protein Assay Kit (Cat. 23225, ThermoFisher Scientific) and denatured at 95°C for 5min with 4x Laemmli Sample Buffer (Cat. 1610747, BioRad). Samples were loaded onto Mini-PROTEAN® TGX^TM^ Gels (Cat. 456-1084, BioRad) and separated by SDS-PAGE for 1.5hrs at 100V in Tris/glycine/SDS running buffer (Cat. 161-0732, BioRad). Proteins were transferred onto 0.45μm nitrocellulose membrane using the Trans-Blot Turbo Transfer System following the manufacturer’s instructions. The membranes were then blocked for 1hr at room temperature using TBST- 5% milk, incubated overnight at 4°C in primary antibody diluted in TBST-5% milk (1:200 ab65796 Kir2.1 (Abcam), 1:500 ab11370 Connexin 43/GJA1(Abcam), 1:2000 C15200203 CRISPR/Cas9 7A9 (Diagenode), 1:1000 ab181602 GAPDH (Abcam)), and incubated at RT for 1hr in secondary antibody diluted in TBST-5% milk (1:4000 ab6721 Goat Anti-Rabbit IgG H&L HRP (Abcam), 1:4000 ab205719 Goat Anti-Mouse IgG H&L HRP (Abcam)). Membranes used to re-probe for multiple proteins were stripped using mild stripping buffer: 1.5% Glycine; 0.1% SDS; 1.0% Tween20; adjusted to pH 2 with HCl). Bands were visualized in with Azure c600 Gel Imaging system (Azure Biosystems) using chemiluminescent substrate Radiance Plus (Cat. AC2103, Azure) and quantified using ImageJ software.

#### Automated Western Blot by Wes

Cells were collected for total protein using Qproteome Mammalian Protein Prep Kit (Cat. 37901, Qiagen). Lysates were quantified for protein expression utilizing the WES system (ProteinSimple) according to the manufacturer’s instructions using a 12-230kDa Separation Module (Cat. SM-W004, ProteinSimple) and either the Anti-Rabbit Detection Module (Cat. DM-001, ProteinSimple) or the Anti-Mouse Detection Module (Cat. DM-002, ProteinSimple). Cell lysates were loaded neat, mixed with 4x Fluorescent Master Mix, and heated at 95°C for 5 minutes. They were blocked with Antibody Diluent (AD), and protein expression was detected with the same primary antibodies listed above. Compared to the standard WB, all antibodies were concentrated by a factor of 20. HRP-conjugated secondary antibodies and chemiluminescent substrate were used directly as indicated as part of the Separation Module. The resulting electropherograms were reviewed and analyzed by ProteinSimple’s WES associated software package, Compass for SW (ProteinSimple), for automated peak detection and quantification. For further details see^61^.

### *t-*SNE Dimension Reduction for Group Visualization

The t-distributed stochastic neighbor embedding (t-SNE) algorithm helps visualize high-dimensional data sets by dimension reduction ^62^. Here, we used MATLAB to produce 2D projections of 19 different measured parameters per sample including: mRNA quantity, action potential duration at 90%, 50% and 30% repolarization (APD80, APD50, APD30), calcium transient duration (CTD90, CTD50, and CTD30) under spontaneous conditions and when paced at 0.5 Hz and 1 Hz. Euclidean measures for the distance between data points and the expression space with perplexity value of 15 were used to visualize clustering of CRISPRi-modified and control sample sets (**Figure 3e**).

### Fluorescence Image Analysis for Levels of Expression

Fluorescence images to visualize expression of the gRNAs-eGFP and dCas9-KRAB-mCherry were adjusted for appropriate LUTs for comparable analysis. To quantify fluorescent images of dCas9 fluorescence time course (**Figure 2**), with mCherry reporter, images were binarized and positive pixel area was quantified as a ratio of the total area. To quantify co-expression ratios in **Figure 4b, f** and **Suppl. Figure 4**, images in each channel (eGFP or mCherry) were adjusted for contrast and binarized using a threshold 0.15-0.21 for the eGFP channel and 0.2-0.3 for the mCherry channel, where all possible values lie between 0-1. Expression ratios were defined by the number of positive pixels in a channel over the total pixels within a field of view or region of interest. For eGFP: *n*_eGFP+_/n_tot_; for mCherry: *n*_mCherry+_/n_tot_; for overlapping eGFP and mCherry: *n_(_*_eGFP+ & mCherry+)_/*n*_eGFP+_, where n_tot_ was the total number of pixels for the camera (512x512). These ratios were calculated and summarized in a violin plot (or box plot) using GraphPad Prism.

### Statistical Analysis

Technical and biological replicates are indicated in the figure legends. All statistical analyses were conducted in Prism Software (GraphPad). One-Way ANOVA followed by post-hoc Tukey tests were used to evaluate significance of Cas9 effects on ion channels compared to eGFP and mCherry controls. Two-Way ANOVA followed by post-hoc Tukey tests were used to evaluate significance between samples with varying pacing frequencies and Dox treatment. Additionally, an unpaired t-test was used to assess the significance between no Dox and Dox treated groups.

## Acknowledgements

This work was supported in part by grants from the National Science Foundation (EFMA1830941 and CBET1705645) and the National Institutes of Health (R01HL144157).

## Author Contributions

JLH and EE designed the experiments. JLH performed all experiments and analyzed the data. WL and YWH built the optical setups used in the macroimaging experiments; WL developed analysis code. YWH and JLH conducted HT all-optical studies and analysis. CC performed image analysis and quantification. EE oversaw the project and secured funding. JLH and EE interpreted the results, wrote and edited the manuscript.

## Data Availability

All data are either shown in the manuscript or made available through github.

## SUPPLEMENTAL INFORMATION

**Supplemental Figure 1.**
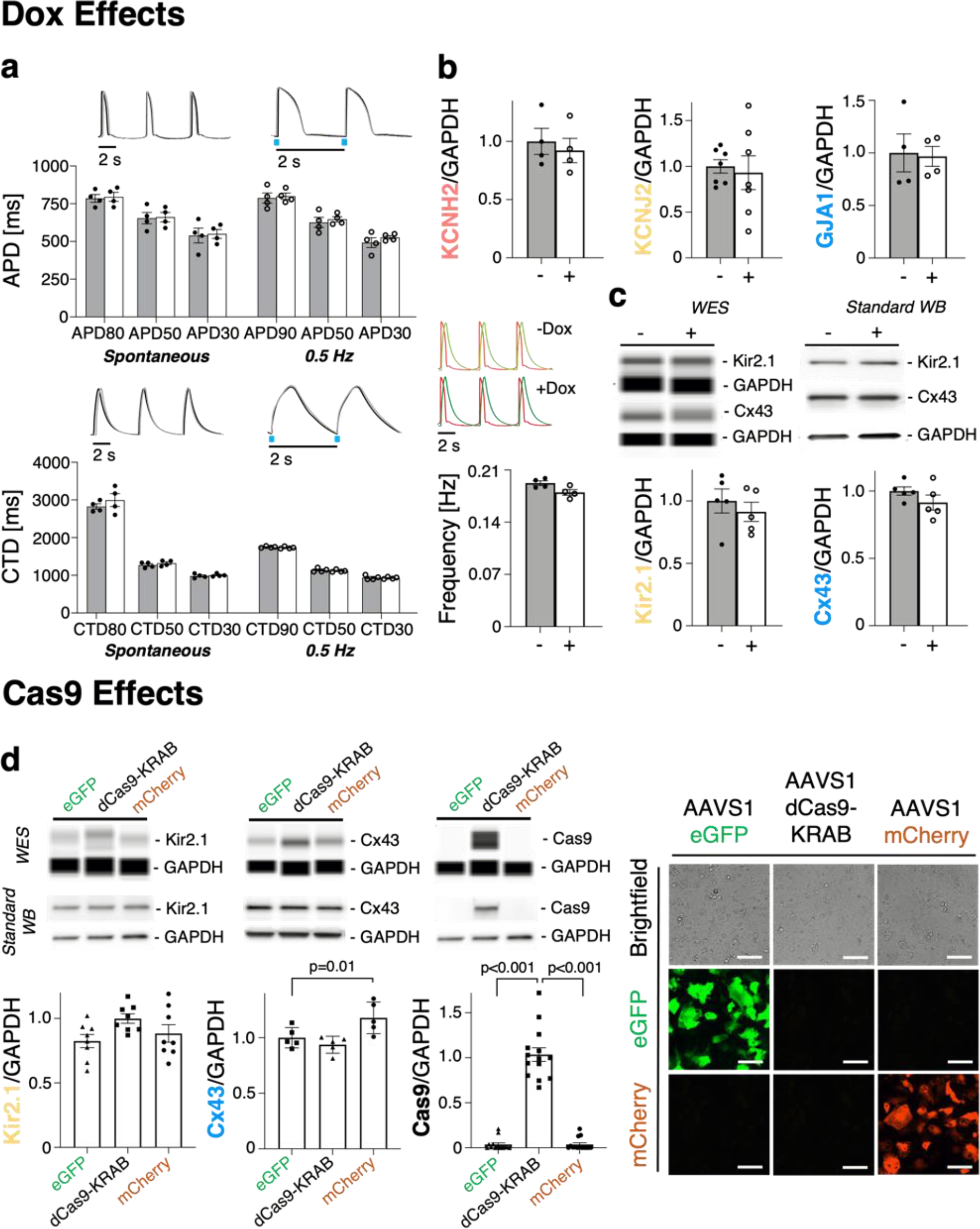
Q**u**antification **of Dox and dCas9-KRAB effects iPSC-CMs on gene expression of ion channels and functional metrics**. (**a-d**) Dox effects: **a,** APD and CTD under spontaneous conditions and 0.5Hz pacing upon 5-day Dox (2 μM) treatment in hiPSC-CMs (n=4 biologically independent samples, two-way ANOVA). Quantification of frequency upon 5-day Dox (2 μM) treatment in hiPSC-CMs (n=4 biologically independent samples; unpaired t-test). Representative voltage (red) and calcium (green) traces. **b,** mRNA quantification of *KCNH2*, *KCNJ2*, and *GJA1* in hiPSC-CMs cultured in Dox (2 μM) for 5 days (n>4 biologically independent samples, n=3 technical replicates; unpaired t-test). **c**, WES (ProteinSimple) and conventional WB of hiPSC-CMs cultured in Dox (2 μM) for 5-days; representative images (WES, n=2 biologically independent samples; WB, n=3 biologically independent samples; unpaired t-test). **d,** dCas9 effects: WES (ProteinSimple) and conventional WB of hiPSC-CMs, expressing eGFP, dCas9-KRAB, and mCherry, with respective plots; representative images (WES, n=2-5 biologically independent samples; WB, n=3-9 biologically independent samples; one-way ANOVA). Data are presented as mean ± SEM. Scale bar 100 *μ*m.

**Supplemental Figure 2.**
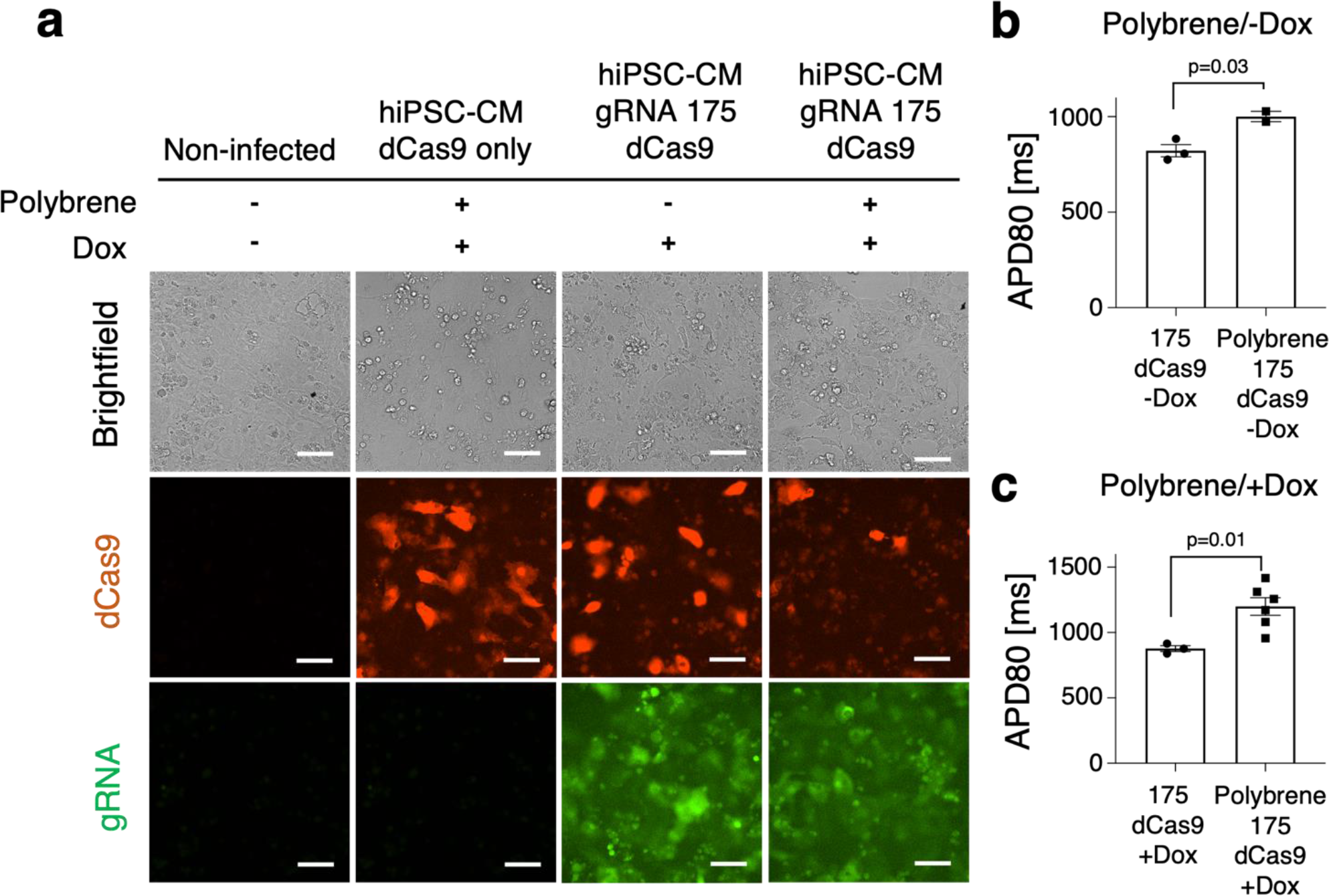
P**o**lybrene **inhibits transfection post-lentivirus transduction**. **a,** Fluorescent images of iPSC-CMs transduced with lentivirus expressing gRNA 175 targeting *KCNH2* with and without Polybrene. Of which Dox-inducible dCas9-KRAB was inserted into the AAVS1 locus. **b,** APD80 of hiPSC-CMs treated with polybrene but no Dox (n>2 biologically independent samples; unpaired t-test). **c,** APD80 of hiPSC-CMs treated with polybrene and Dox (n>3 biologically independent samples; unpaired t-test). Data are presented as mean ± S.E.M. Scale bar 100 *μ*m.

**Supplemental Figure 3.**
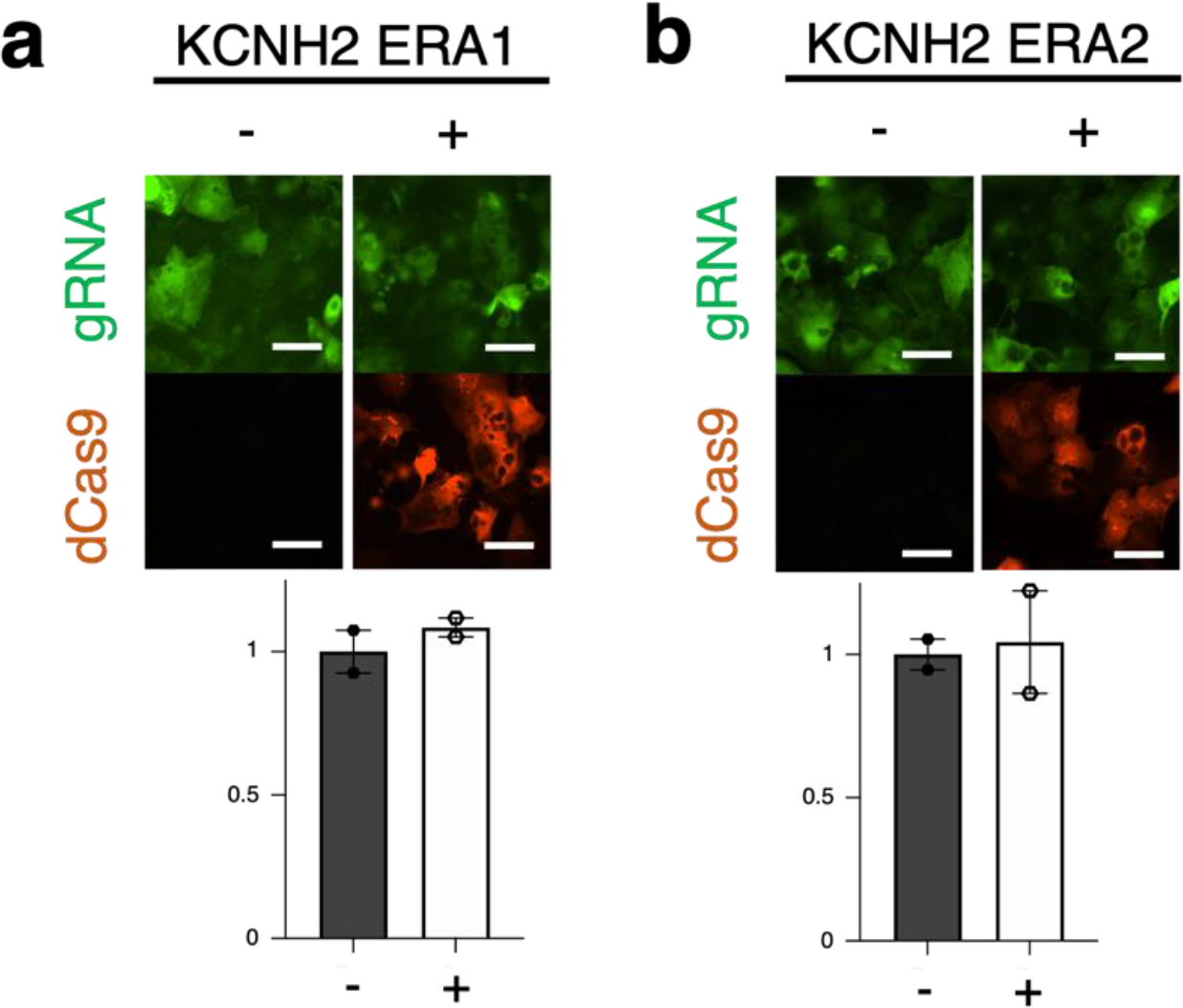
Additional gRNA’s tested for *KCNH2* knockdown efficiency with *KCNH2* **ERA1 and *KCNH2* ERA2.** Data are presented as mean ± S.E.M. Scale bar 100 *μ*m.

**Supplemental Figure 4.**
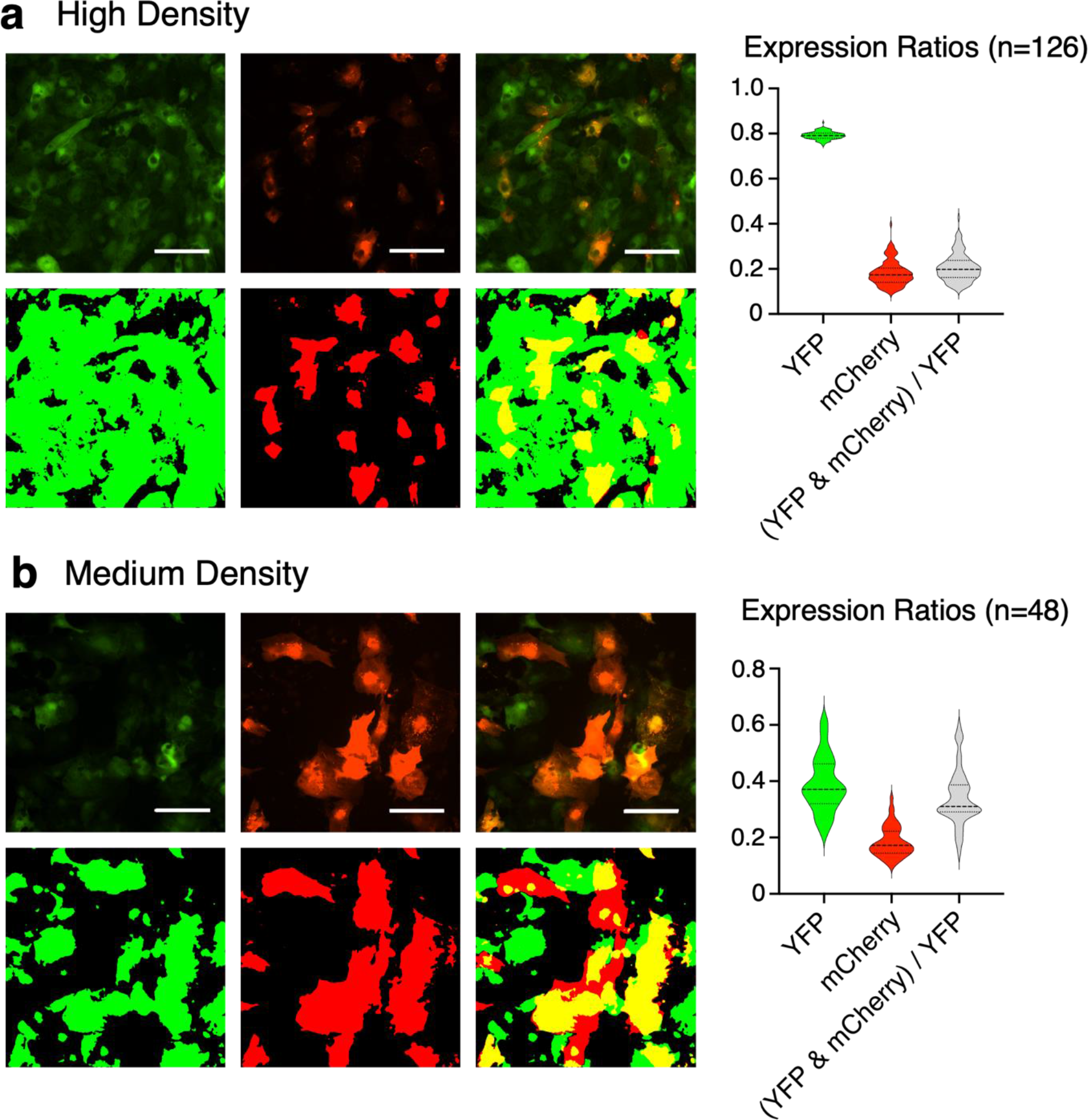
Q**u**antification **of fluorescence expression ratios of mCherry tagged dCas9- KRAB and eGFP tagged gRNA**. **a,** Images of high density groups (n=126 independent ROI), scale bar 100 *μ*m, and medium density groups (n=48 independent ROI), scale bar 50 *μ*m, were analyzed for pixel quantification to calculate regions of overlap in expression of dCas9-KRAB and gRNA. **b,** Plots of expression ratios quantified.

**Supplemental Figure 5.**
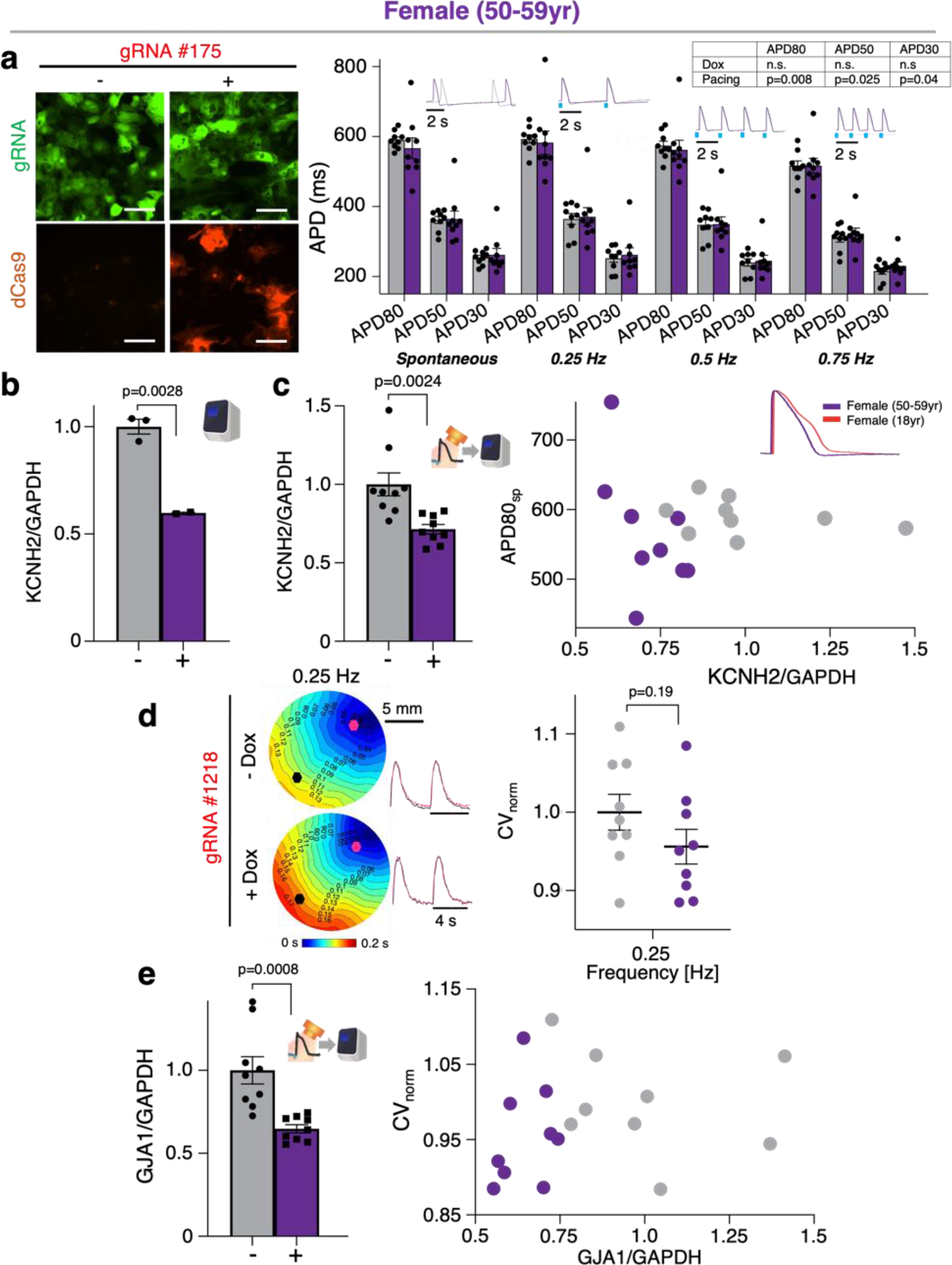
C**R**ISPRi **modulation of female diversity line**. **a,** Fluorescence expression of dCas9-mCherry and gRNA 175 tagged with eGFP upon 5-days Dox (2 μM) treatment in female (50-59yr) hiPSC-CMs; representative images. Functional changes in APD in Dox (2 μM) treated samples expressing Dox-inducible dCas9-KRAB and gRNA targeting *KCNH2* (n=9 biologically independent samples; two-way ANOVA). **b,** qPCR of CRISPRi knockdown of *KCNH2* with gRNA 175 in female hiPSC-CMs (n>2 biologically independent samples, n=3 technical replicates; unpaired t-test). **c,** qPCR analysis of *KCNH2* knockdown post-staining and functional experiments (n=9 biologically independent samples, n=3 technical replicates; unpaired t-test). Plot correlating the effects of relative *KCNH2* mRNA and spontaneous APD80 (n=9 biologically independent samples). **d,** Activation maps of female hiPSC-CMs CRISPRi *GJA1* knockdown and gRNA 1218 at 0.25 Hz pacing; representative images (n=9 biologically independent samples). Conduction velocity changes in Dox (2 μM) treated samples at 0.25 Hz pacing (n=9 biologically independent samples, unpaired t-test). **e,** qPCR of CRISPRi knockdown of *GJA1* with gRNA 1218 post-staining and functional experiments (n=9 biologically independent samples, n=3 technical replicates; unpaired t-test). Plot correlating relative *GJA1* mRNA levels with CV at 1 sec (n=9 biologically independent samples). Data are presented as mean ± S.E.M. Scale bar (100 *μ*m).

**Supplemental Figure 6.**
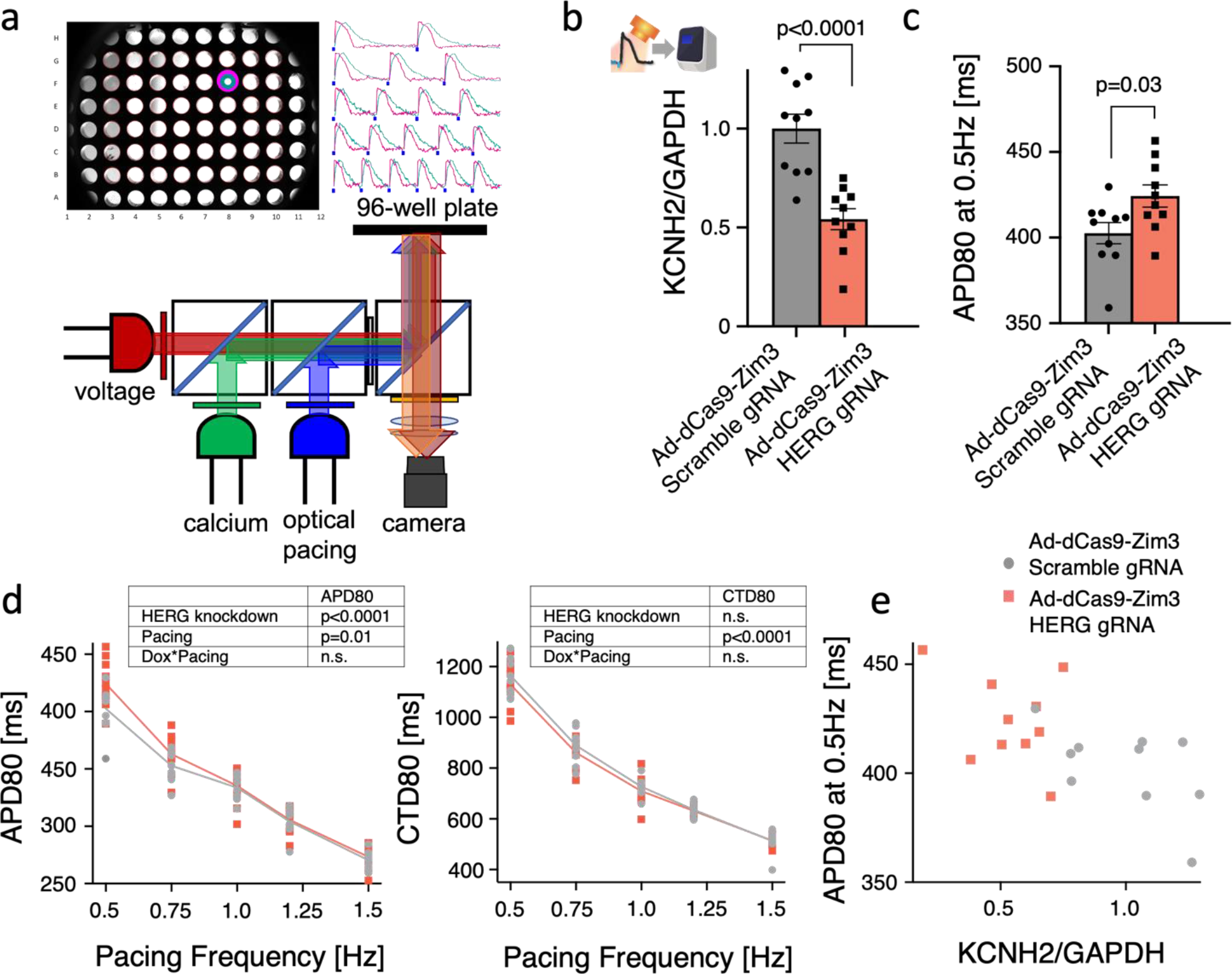
S**u**pplementary **High-throughput (HT) studies for synchronous, multi-well measurement**. **a**, Schematic illustrating optical system capable of simultaneous optical pacing and voltage and calcium imaging of an entire 96-well plate. HT, all-optical electrophysiology studies were conducted at 35°C. **b**, qPCR of dCas9-Zim3 knockdown of *KCNH2* post-staining and functional experiments (n=10 biologically independent samples, n=3 technical replicates; unpaired t-test). **c**, Functional changes in APD80 at 0.5 Hz pacing upon *KCNH2* inhibition by dCas9-Zim3 (n=10 biologically independent samples; unpaired t-test). **d**, Restitution plot of APD80 and CTD80 upon *KCNH2* knockdown (n=10 biologically independent samples; two-way ANOVA). **e**, Plot correlating relative *KCNH2* mRNA levels with APD80 at 0.5 Hz pacing (n=10 biologically independent samples). Data are presented as mean ± S.E.M.

**Supplemental Table 1.** **Cell lines tested.**

| CELL TYPE | SEX | Tissue Source | AGE | SOURCE | IDENTIFIER |
| --- | --- | --- | --- | --- | --- |
| iCell<sup>2</sup> | Female | Fibroblast | 18yr | FujiFilm/Cellular Dynamics | 01434 |
| iCell Diversity Panel | Female | N/A | 50-59yr | FujiFilm/Cellular Dynamics | iPSC-CM-1X-01064 |
| iCell Diversity Panel | Male | N/A | 50-59yr | FujiFilm/Cellular Dynamics | iPSC-CM-1X-01063 |

**Supplemental Table 2.**
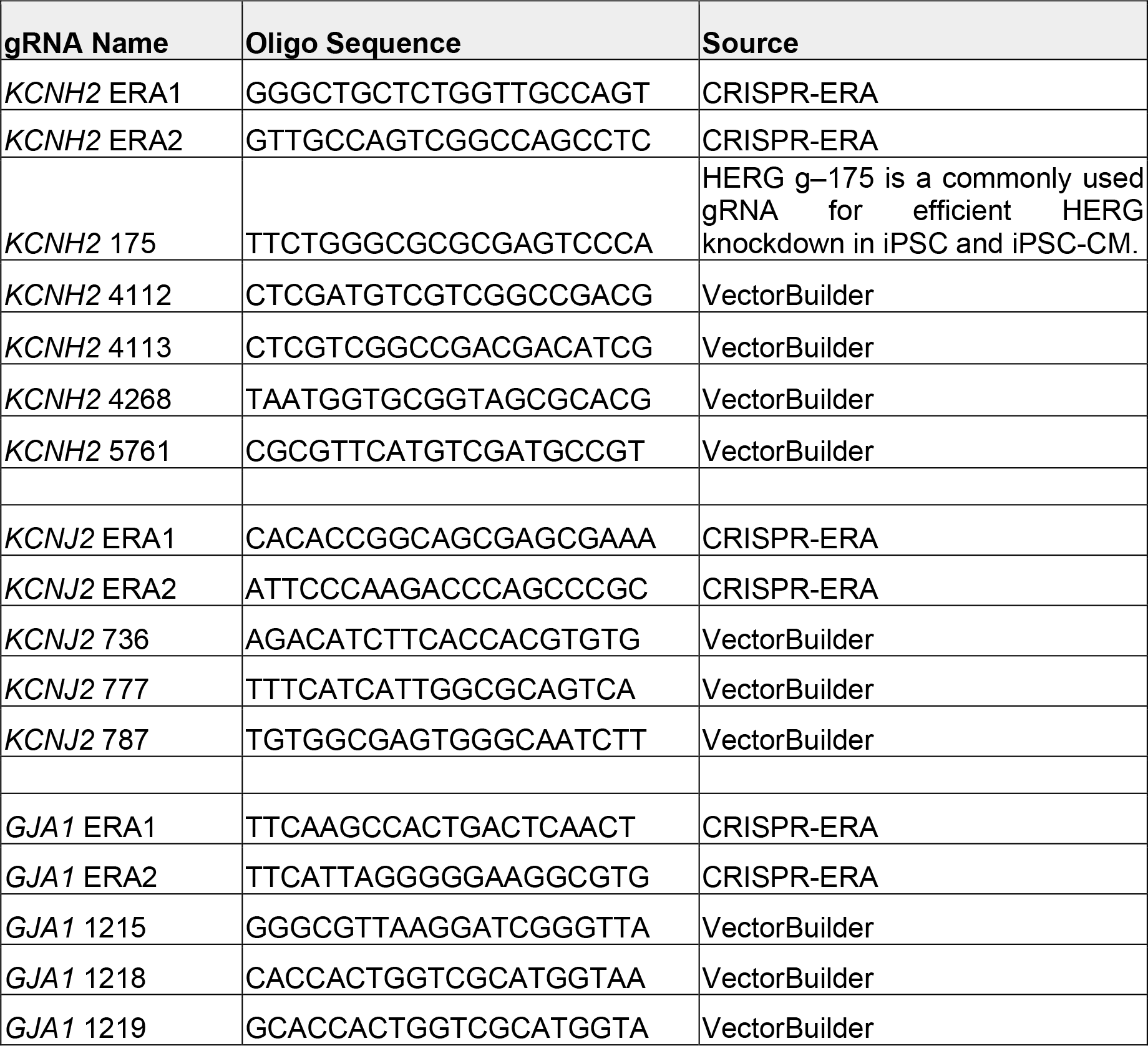
**gRNA oligo sequences tested, related to Methods**.

**Supplemental Table 3.**
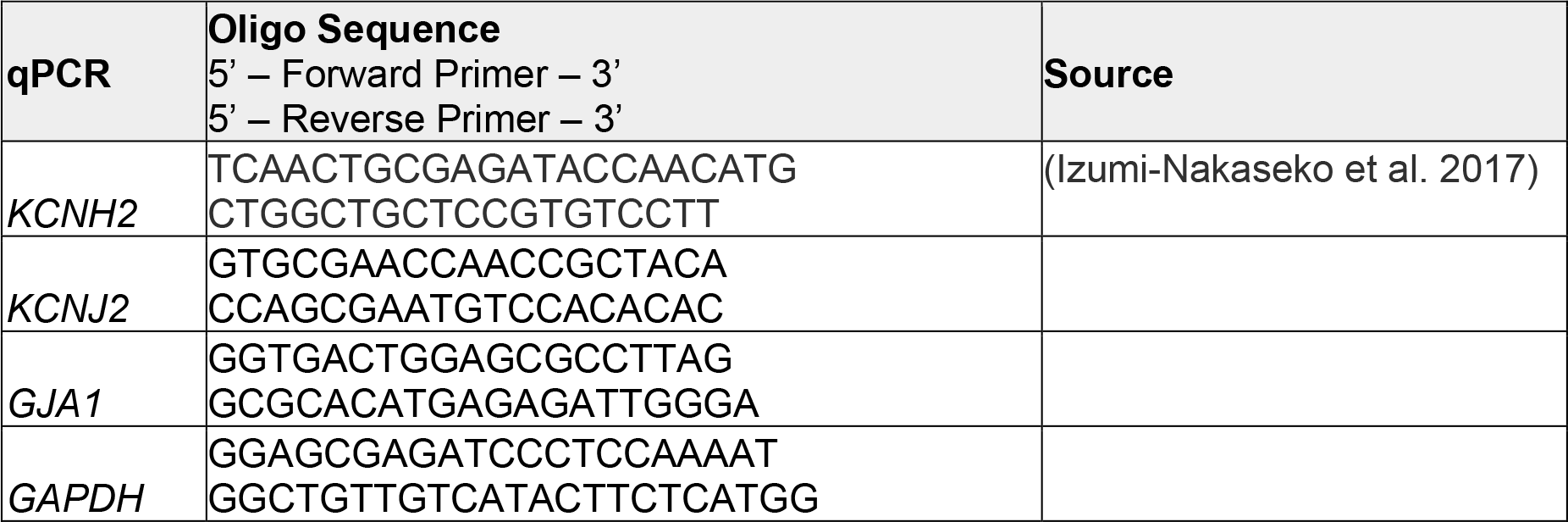
**Primers used for qPCR Analysis**.

